# Tissue-specific effects of dietary protein on cellular senescence are mediated by branched-chain amino acids

**DOI:** 10.1101/2025.01.13.632607

**Authors:** Mariah F. Calubag, Ismail Ademi, Isaac Grunow, Lucia Breuer, Reji Babygirija, Penelope Lialios, Sandra Le, Dennis Minton, Michelle M. Sonsalla, Julia Illiano, Bailey A. Knopf, Fan Xiao, Adam R. Konopka, David A. Harris, Dudley W. Lamming

## Abstract

Dietary protein is a key regulator of healthy aging in both mice and humans. In mice, reducing dietary levels of the branched-chain amino acids (BCAAs) recapitulates many of the benefits of a low protein diet; BCAA-restricted diets extend lifespan, reduce frailty, and improve metabolic health, while BCAA supplementation shortens lifespan, promotes obesity, and impairs glycemic control. Recently, high protein diets have been shown to promote cellular senescence, a hallmark of aging implicated in many age-related diseases, in the liver of mice. Here, we test the hypothesis that the effects of high protein diets on metabolic health and on cell senescence are mediated by BCAAs. We find that reducing dietary levels of BCAAs protects male and female mice from the negative metabolic consequences of both normal and high protein diets. Further, we identify tissue-specific effects of BCAAs on cellular senescence, with restriction of all three BCAAs – but not individual BCAAs – protecting from hepatic cellular senescence while potentiating cell senescence in white adipose tissue. We find that the effects of BCAAs on hepatic cellular senescence are cell-autonomous, with lower levels of BCAAs protecting cultured cells from antimycin-A induced senescence. Our results demonstrate a direct effect of a specific dietary component on a hallmark of aging and suggest that cellular senescence may be highly susceptible to dietary interventions.

## Introduction

As the global population ages, there is growing interest – and urgency – to identify effective and affordable interventions to promote healthy aging ^1^. One of the 12 hallmarks of aging ^2^, cellular senescence (CS) has attracted significant attention as a potential target for therapeutic intervention as the accumulation of senescent cells and their associated mix of cytokines, chemokines, and other factors, collectively known as the senescence-associated secretory phenotype (SASP), is linked to a broad array of age-related diseases ^3–18^.

While, dietary interventions are appealing, calorie restriction (CR), the gold standard for aging interventions, is difficult for many ^19^. Recently dietary composition – especially dietary protein – has been shown to have strong effects on healthy aging ^20^. While more protein is typically thought of as beneficial, higher protein intake is associated with increased mortality and age-related diseases in humans ^21,22^. Conversely, protein restriction (PR) promotes metabolic health in both humans and rodents, and extends rodent lifespan ^23–29^. It was recently shown that as dietary protein increases, hepatic CS increases ^30^.

While the mechanism by which dietary protein promotes CS is unknown, many of the metabolic benefits of PR are mediated by reduced dietary levels of the three branched-chain amino acids (BCAAs; leucine, isoleucine, and valine). BCAAs are powerful regulators of healthy aging, with restriction of all three BCAAs or isoleucine alone improving metabolic health, reducing frailty, and extending lifespan ^26,29,31^. In contrast, BCAA supplementation promotes obesity and insulin resistance, and reduces lifespan ^32,33^. An accumulating set of data suggests that BCAAs promote senescence, with BCAA supplementation promoting senescence in cell culture experiments ^34^ as well as *in vivo* in mice ^35^. Genetic manipulation of BCAA catabolism in cells and mice likewise is consistent with BCAAs promotes CS ^36,37^.

Thus, it is possible that dietary protein promotes CS via the BCAAs. If this hypothesis is correct, several molecular mechanisms could contribute to BCAA-mediated CS. Protein in general, and the BCAAs in particular, are potent agonists of the mechanistic Target of Rapamycin Complex 1 (mTORC1), a highly conserved protein kinase that, when inhibited, extends the lifespan of mice ^38,39^. Protein restriction and BCAA restriction both reduce mTORC1 activity in mice ^25,40^, and mTORC1 promotes the SASP; rapamycin, which extends mouse lifespan by inhibiting mTORC1, inhibits the SASP ^41^. Additionally, both PR and BCAA restriction induce fibroblast growth factor 21 (FGF21), a hormone that extends lifespan when overexpressed and regulates mTORC1 in multiple tissues ^26,42–45^. *In vitro*, knockdown of FGF21 promotes accumulation of senescent cells ^46^ whereas FGF21 overexpression and administration has been shown to protect against CS ^46,47^.

In this study, we tested the hypothesis that the effects of dietary protein on CS are mediated by the BCAAs. We found that restriction of the dietary levels of BCAAs protects male and female mice from the negative metabolic consequences of both normal and high protein diets, and that the dietary levels of BCAAs – but not protein – are associated with hepatic CS and SASP gene expression.

Further, we discovered tissue-specific effects of BCAAs on cellular senescence, with restriction of all three BCAAs – but not individual BCAAs – protecting from hepatic cellular senescence while potentiating cell senescence in white adipose tissue. Despite previous literature, we found that this tissue-specific effect is independent of the FGF21/mTORC1 axis, but rather mediated by an effect of BCAAs on mitochondrial function. We found that the effect of BCAAs on hepatic cellular senescence are cell-autonomous, with lower levels of BCAAs protecting cultured cells from antimycin-A induced senescence. We conclude that dietary BCAAs can promote metabolic health even in the context of higher protein diets, and that BCAAs are a dietary component that affects senescent cell accumulation in a tissue-specific manner.

## Materials and Methods

### Animal care, housing and diet

All procedures were performed in conformance with institutional guidelines and were approved by the Institutional Animal Care and Use Committee (IACUC) of the William S. Middleton Memorial Veterans Hospital and the University of Wisconsin-Madison IACUC. Male C57BL/6J mice were purchased from The Jackson Laboratory (Bar Harbor, ME, USA) at 11 weeks of age. p16-3MR reporter mice, in which the p16^INK4a^ promoter drives expression of the 3MR fusion protein, ^48^ were received at 13 weeks of age from the Harris and Ricke labs at UW-Madison, which were generously gifted on the C57BL/6J background by Dr. Judith Campisi. All mice were acclimated to the animal research facility for at least one week before entering studies. All animals were housed in static microisolator cages in a specific pathogen-free mouse facility with a 12:12 h light– dark cycle, maintained at approximately 22°C.

Mice were fed amino acid defined diets with varying levels of protein and BCAAs (full diet compositions are provided in **Table S1**; Inotiv, Madison, WI, USA). Diets were started at either 12 weeks of age or 16 months of age in the C57BL/6J mice or at 14 weeks of age in the p16-3MR mice, and continued for at least 16 weeks. 8-week-old male C57BL/6J mice were placed on a high-fat high sucrose western diet (WD; TD.160186) or a WD with restricted BCAAs (WD-BR; TD.160188) for 45 weeks.

### Metabolic Phenotyping

Glucose, insulin and alanine tolerance tests were performed by fasting all mice for 4 hours or overnight (∼16 hours) and then injecting either glucose (1g/kg), insulin (0.75U/kg) or pyruvate (2g/kg) intraperitoneally ^49,50^.

Blood glucose levels were determined at the indicated times using a Bayer Contour blood glucose meter (Bayer, Leverkusen, Germany) and test strips. Mouse body composition was determined using an EchoMRI Body Composition Analyzer. For assay of multiple metabolic parameters (O2, CO2, food consumption, and activity tracking), mice were acclimatized to housing in a Columbus Instruments Oxymax/CLAMS-HC metabolic chamber system for approximately 24 hours, and data from a continuous 24-hour period was then recorded and analyzed.

### In Vivo Bioluminsescence

*In vivo* luminescence assessment was performed by the Small Animal Imaging & Radiotherapy Facility (SAIRF). p16-3MR reporter mice were injected intraperitoneally with 15 μg of RediJect Coelenterazine h Bioluminescent Substrate (Fischer Scientific; Catalog No.50-209-9325). 15 minutes later, the mice were anesthetized with isofluorane, and luminescence was measured with a Revvity IVIS Spectrum *in vivo* imaging system (Revvity Health Sciences; 1 minute medium binning).

### Collection of tissues for molecular and histological analysis

Mice were euthanized in the fasted or fed state at the indicated age. Mice euthanized in the fed state were fasted overnight starting the day prior to sacrifice; in the morning, mice were refed for 3 hours and then sacrificed 3 hours later. Following blood collection via submandibular bleeding, mice were euthanized by cervical dislocation and tissues were rapidly collected, weighed, and snap frozen in liquid nitrogen. A portion of the liver was fixed in 10% formalin for 4 hours, transferred to sucrose for 24 hours and then embedded in Tissue-Tek Optimal Cutting Temperature (OCT) compound, and then cryosectioned and stained for SA-β-gal activity. Images of the liver were taken using an EVOS microscope (Thermo Fisher Scientific Inc., Waltham, MA, USA) at a magnification of 40X as previously described ^31,51,52^. For quantification of lipid droplet size, six independent fields were obtained for each tissue from each mouse and quantified using ImageJ (NIH, Bethesda, MD, USA).

### Cell Culture and treatment

AML12 cells were obtained from ATCC. AML12 cells were cultured in Gibco Dulbecco’s modified Eagle’s medium with F12 (DMEM/F12) without Amino Acids, Glucose, L-Glutamine, Sodium Bicarbonate, HEPES, Sodium Pyruvate, Hypoxanthine, Thymidine, Phenol Red (Powder) (D9807-10; US Biological Life Sciences, Waltham, MA, USA) supplemented with 10% fetal bovine serum (FBS) (12306C; Life Technologies, Carlsbad, CA, USA) and 1% Penicillin/Streptomycin (15140122; Gibco, Billings, MT, USA). Amino acids and glucose were added to create custom DMEM lacking BCAAs. See Table **S5** for amino acid and complete cell culture composition and catalog numbers. CS was induced in AML12 cells by treating them with 1uM Antimycin A (A8674-25MG; Sigma, St. Louis, MO, USA) for 14-21 days prior to collection.

### Quantitative real-time PCR

RNA was extracted from liver or inguinal white adipose tissue (iWAT) using TRI Reagent according to the manufacturer’s protocol (Sigma-Aldrich). The concentration and purity of RNA were determined by absorbance at 260/280 nm using Nanodrop (Thermo Fisher Scientific). 1 μg of RNA was used to generate cDNA (Superscript III; Invitrogen, Carlsbad, CA, USA). Oligo dT primers and primers for real-time PCR were obtained from Integrated DNA Technologies (IDT, Coralville, IA, USA); sequences are in **Table S2**. Reactions were run on an StepOne Plus machine (Applied Biosystems, Foster City, CA, USA) with Sybr Green PCR Master Mix (Invitrogen). Actin was used to normalize the results from gene-specific reactions.

### Immunoblotting

Animals used for Western blotting were sacrificed following an overnight fast and 3-hour refeed. Tissue samples from muscle were lysed in cold RIPA buffer supplemented with phosphatase inhibitor and protease inhibitor cocktail tablets (Thermo Fisher Scientific, Waltham, MA, USA) as previously described ^29,53^ using a FastPrep 24 (M.P. Biomedicals, Santa Ana, CA, USA) with screw cap microcentrifuge tubes (822-S) from (Dot Scientific, Burton, MI) and ceramic oxide bulk beads (10158-552) from VWR (Radnor, PA, USA). Protein lysates were then centrifuged at 13,300 rpm for 10 min and the supernatant was collected. Protein concentration was determined by Bradford (Pierce Biotechnology, Waltham, MA, USA). 20 μg protein was separated by SDS–PAGE (sodium dodecyl sulfate–polyacrylamide gel electrophoresis) on 16% resolving gels (Thermo Fisher Scientific, Waltham, MA, USA) and transferred to PVDF membrane (EMD Millipore, Burlington, MA, USA). pT389-S6K1 (9234), S6K1 (9202), pS240/244-S6 (2215), S6 (2217), p-Thr37/46 4E-BP1 (2855), and 4E-BP1 (9644) were purchased from Cell Signaling Technologies (CST, Danvers, MA, USA) and used at a dilution of 1:1000. Imaging was performed using a Bio-Rad Chemidoc MP imaging station (Bio-Rad, Hercules, CA, USA). Quantification was performed by densitometry using NIH ImageJ software.

### Senescence-Associated B-Galactosidase Staining

Enhanced lysosomal biogenesis, a common feature of cellular senescence, can be detected by measuring β-galactosidase (lysosomal hydrolase) activity at pH 6.0 ^54^. This senescence-associated β-galactosidase (SA-β-gal) activity appears to be restricted to senescent cells at a low pH ^55,56^. SA-β-gal activity is shared by almost all senescent cells, but particular care should be taken when detecting this in cultured cells because high confluency and contact inhibition can increase SA-β-gal activity ^55,57^. Fresh tissue from mice was collected and fixed in 10% neutral buffered formalin (NBF) on ice for 3-4 hr. Tissues were then transferred to 30% sucrose at 4°C for 24 hours before being embedded in O.C.T. compound in a cryomold and stored at -80°C. Prior to cryosectioning, tissues were equilibrated at -20°C and then cryosectioned into 5-7 µm sections before being attached to Superfrost Plus slides. Fresh SA-β-gal staining solution at a pH of 6 was prepared as previously described. Tissue slides were stained in SA-β-gal staining solution for 18-24 hrs at 37°C in a non-CO2 incubator before being rinsed three times with PBS. To prevent crystal formation a parafilm Coplin jar was used to prevent evaporative loss of staining solution. Stained sections were imaged using the EVOS microscope (Thermo Fisher Scientific Inc., Waltham, MA, USA) at a magnification of 40X as previously described. The percent of SA-βgal-positive area for each sample will be quantified using ImageJ.

### ELISA assays and kits

Blood plasma for FGF21 was obtained at week 15 in the fasted state and from blood collected at time of euthanasia in the refed state. Blood FGF21 levels were assayed by a mouse/rat FGF-21 quantikine ELISA kit (MF2100) from R&D Systems (Minneapolis, MN, USA).

### Statistics

Data are presented as the mean ± SEM unless otherwise specified. Statistical analyses were performed using one-way or two-way ANOVA followed by Tukey–Kramer post hoc test, as specified in the figure legends. Outliers were excluded using the Robust Regression Outlier Test (ROUT) in Graphpad Prism (v10), Q=1%, and are indicated by an asterik(*) and blue colored font in the Source Data. Other statistical details are available in the figure legends. Energy expenditure differences were detected using analysis of covariance (ANCOVA). ANCOVA analysis assumes a linear relationship between the variables and their covariates. If the slope is equal between groups, then the regression lines are parallel, and elevation is then tested to determine any differences (i.e., if slopes are statistically significantly different, elevation will not be determined). In all figures, n represents the number of biologically independent animals. Sample sizes were chosen based on our previously published experimental results with the effects of dietary interventions ^26,31,49,52,58^. Data distribution was assumed to be normal, but this was not formally tested.

### Randomization

All studies were performed on animals or on tissues collected from animals. Young animals of each sex and strain were randomized into groups of equivalent weight, housed 2-3 animals per cage, before the beginning of the *in vivo* studies. Aged animals were randomized into groups of equivalent weight, housed 2-3 animals per cage, when possible.

## Results

### Restricting BCAAs improves the metabolic health and increases energy expenditure of mice eating normal and high protein diets

We began our study by designing a series of diets in which the total amount of calories derived from amino acids (AAs) was either 7% (LP, Low Protein), 21% (CP, Control Protein), or 36% (HP, High Protein). These diets were isocaloric. Dietary fat was held constant (19%) and the levels of carbohydrates adjusted in order maintain equivalent caloric density. We further designed CP and HP diets in which the levels of all three BCAAs were reduced to the level of BCAAs found in the LP diet (CP-BR and HP-BR, respectively); in these diets, the reduction in BCAAs was balanced by a proportional increase in non-essential AAs, keeping the percentage of calories derived from AAs constant. The full composition of these diets is summarized in **Table S1**.

We randomized 12-week-old C57BL/6J male mice to these diets, following them longitudinally for 16 weeks (**Fig. 1A**). As we anticipated based on our previous studies, we found that mice consuming diets low in BCAAs (i.e., LP, CP-BR, HP-BR) gained significantly less weight and fat mass than CP- and HP-fed mice, with an overall reduction in adiposity (**Supplementary Figs. S1A-H**). Importantly, this weight reduction was not due to decreased calorie intake; mice consuming the LP, CP-BR, and HP-BR diets consumed more calories than CP and HP-fed mice (**Supplementary Figs. S1I-L**). Due to this increase, animals consuming the CP-BR and HP-BR diets had an overall increase in calories derived from AAs, even while BCAAs were effectively restricted (**Supplementary Figs. S1M-P**).

**Figure 1:**
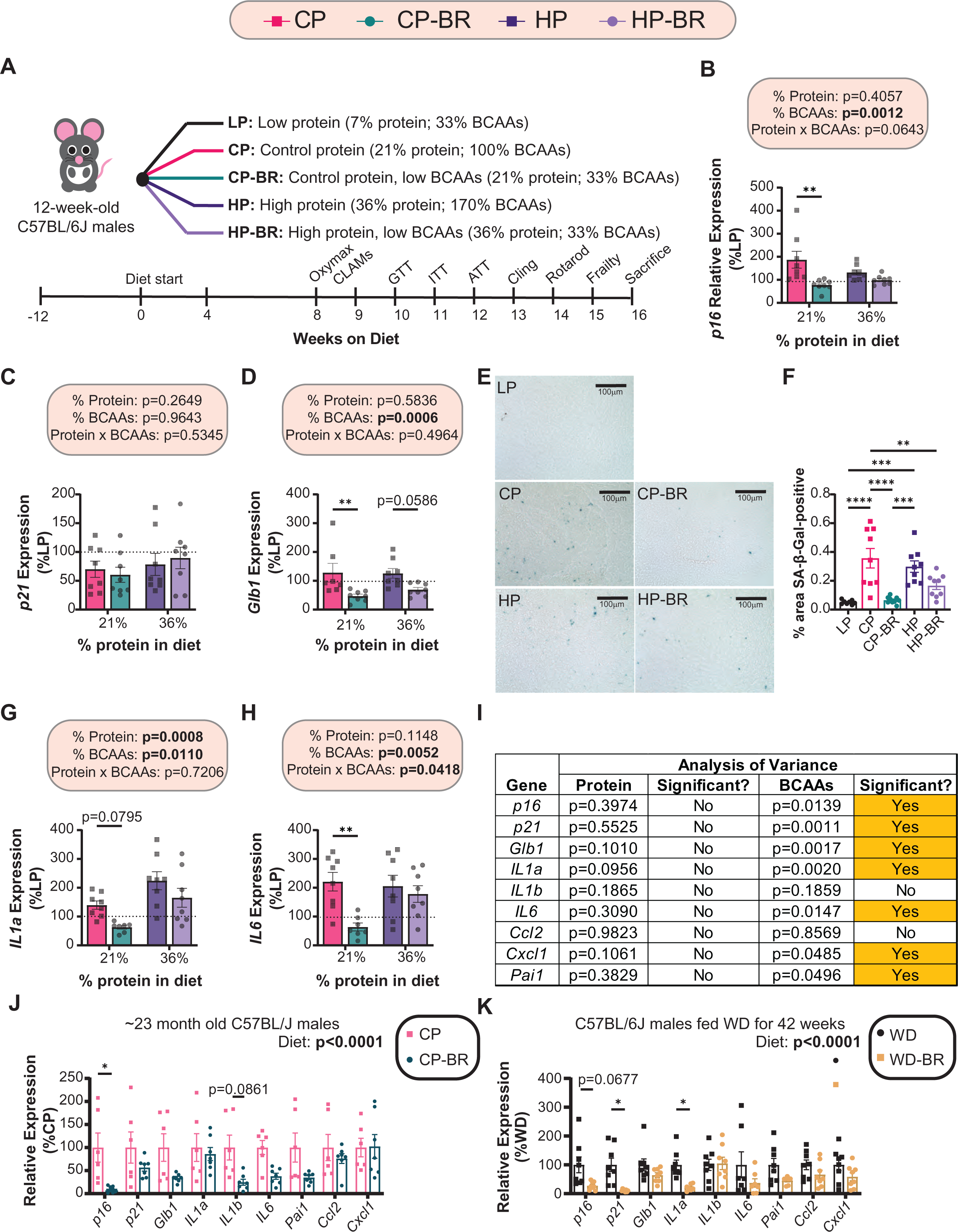
Diets low in BCAAs protect C57BL/6J male mice from the accumulation of senescent cells in the liver. (A) Experimental design. (B-D) Hepatic mRNA expression of *p16, p21* and *Glb1* in mice fed the indicated diets. (E-F) Hepatic SA-βGal staining at 40X magnification (scale bar = 100 µm) (E) with quantification of SA-βGal-positive cells (F). (G-H) Hepatic mRNA expression of *IL1a* and *IL6* in mice fed the indicated diets. (I) Multiple linear regression (MLR) analysis to determine contribution of protein versus BCAAs to hepatic senescence gene expression (*p16, p21, Glb1, IL1a, IL1b, IL6, Ccl2, Cxcl1* and *Pai1*). (J) Hepatic mRNA expression of senescence genes in male C57BL/6J mice on CP and CP-BR diets from 16 months of age until 23 months of age. (K) Hepatic mRNA expression of senescence genes in ∼48 week old male mice after a ∼42 weeks consuming a WD-BR diet. (B-D, G-H) n=7-8 mice/group (100% = average expression of the indicated gene in the liver of LP-fed mice). The overall effect of protein, BCAAs, and the interaction represent the p-value from a two-way ANOVA; *p<0.05, Sidak’s test post 2-way ANOVA. (F) n=8-9 mice/group; *p<0.05, Tukey test post ANOVA. (I) Statistics for the p-value are from a MLR analysis to determine the contribution of protein versus BCAAs from the senescence data set. (J-K) n=6-8 mice/group. The overall effect of diet represents the p-value from a two-way ANOVA; *p<0.05, Sidak’s test post 2-way ANOVA conducted separately for each gene. Data represented as mean ± SEM.

We have previously observed that the paradoxical decreased weight gain of BCAA-restricted mice is associated with increased energy expenditure ^26,31^. Using metabolic chambers, we determined energy expenditure via indirect calorimetry while also assessing food consumption, activity, and fuel source utilization. As we anticipated, mice with reduced levels of BCAAs had increased energy expenditure relative to the respective BCAA-replete diet (**Supplementary Figs. S2A-E**). We also noted a tendency for increased spontaneous activity in animals with lower levels of BCAAs that reached statistical significance in some cases (**Supplementary Fig. S2F**).

Increased energy expenditure in LP-fed mice is mediated by FGF21, which is induced by PR and stimulates the beiging of inguinal white adipose tissue (iWAT) and the activation of brown adipose tissue (BAT) by stimulating sympathetic nerve activity ^59–61^. We observed that BCAA restriction increased plasma FGF21 in the context of the CP (21% protein) diet, but not the HP diet (36% protein diet) (**Supplementary Figs. S2G-H**). We observed that thermogenic genes were upregulated in the inguinal white adipose tissue (iWAT) in response to an LP diet, but not response to BCAA restriction; conversely, in BAT, we observed an BCAA-restriction induction of several thermogenic genes (**Supplementary Figs. S2I-J**). Mice consuming diets low in BCAAs have a higher respiratory exchange ratio (RER), suggesting an increase in carbohydrate/protein substrate utilization (**Supplementary Figs. 2K-M**). This data is consistent with BCAA restriction increasing energy expenditure via FGF21-induced activation of thermogenesis.

We also assessed glucose homeostasis by performing glucose, insulin, and alanine tolerance tests. Mice consuming LP and low BCAA diets had improved glucose and pyruvate tolerance and lower fasting blood glucose relative to mice consuming CP and HP diets; however, insulin sensitivity was improved only in LP-fed mice (**Supplementary Figs. S3A-G**).

### LP- and CP-BR-fed animals decreased hepatic cellular senescence

As described above, we hypothesized that the metabolic benefits of low BCAA diets are in part through altered CS. We therefore examined the mRNA expression of common senescence and SASP genes. We observed that BCAA restriction reduced the expression of the CS markers *p16* and *Glb1*, while the expression of *p21* was not significantly altered (**Figs. 1B-D**). We also performed hepatic SA-β-gal staining and livers from mice fed diets with low levels of BCAAs (LP, CP-BR and HP-BR) had lower SA-β-gal-positive staining than livers from CP and HP-fed mice, matching with the *Glb1* gene expression (**Fig. 1D-F**). BCAA restriction also had strong effects on the expression of multiple SASP genes, reducing expression of *Il1a, Il6, Ccl2,* and *Cxcl1*, while not affecting the expression of several other SASP genes (**Figs. 1G-H and Supplementary Figs. 4A-D**).

The effects of BCAA restriction were generally stronger in the context of the CP diet, while BCAA restriction in the context of the HP diet either had no effect or did not reach statistical significance. To better understand the roles of BCAAs and protein in the regulation of these CS and SASP genes, we analyzed the mRNA expression using a Multiple Linear Regression (MLR) approach with gene expression as the dependent variable and the level of either dietary protein or BCAAs as the independent variables. We found that dietary BCAAs substantially contributed to the expression of most of the CS and SASP genes we examined, while dietary protein level did not (**Fig. 1I**).

We examined the effects of BCAA restriction on CS and the SASP in other contexts to determine the general applicability of these findings. We found that there was an overall beneficial effect of BCAA restriction on CS and SASP gene expression in the livers of aged C57BL/6J male mice, with restriction of BCAAs significantly reducing expression of *p16* (**Fig. I-J**). We further found that BCAA restriction had an overall beneficial effect on CS and SASP gene expression in the livers of mice fed a high-fat, high-sucrose Western diet, significantly reducing expression of *p16*, *p21*, and *Il1a* (**Fig. 1K**).

### Low BCAA diets display a tissue-specific effect on adipose tissue

We repeated this study in p16-3MR mice. Overall, the effects of protein and BCAAs on body weight, food intake, and glucose regulation in p16-3MR males was similar to what we observed in C57BL/6J males (**Supplementary Fig. 5**).

At 20-22 weeks of age, we analyzed p16 expression *in vivo* using a bioluminescence assay. Surprisingly, we found no significant difference between mice fed any of the diets on whole-body total flux, which represents p16 expression (**Figs. 2A-B**). When we plotted the total flux against either protein or BCAA consumption, we found that there was a statistically significant relationship between BCAAs and flux; surprisingly though, the slope in response to BCAAs was negative, suggesting a negative association between BCAAs and CS (**Figs. 2C-D**).

**Figure 2:**
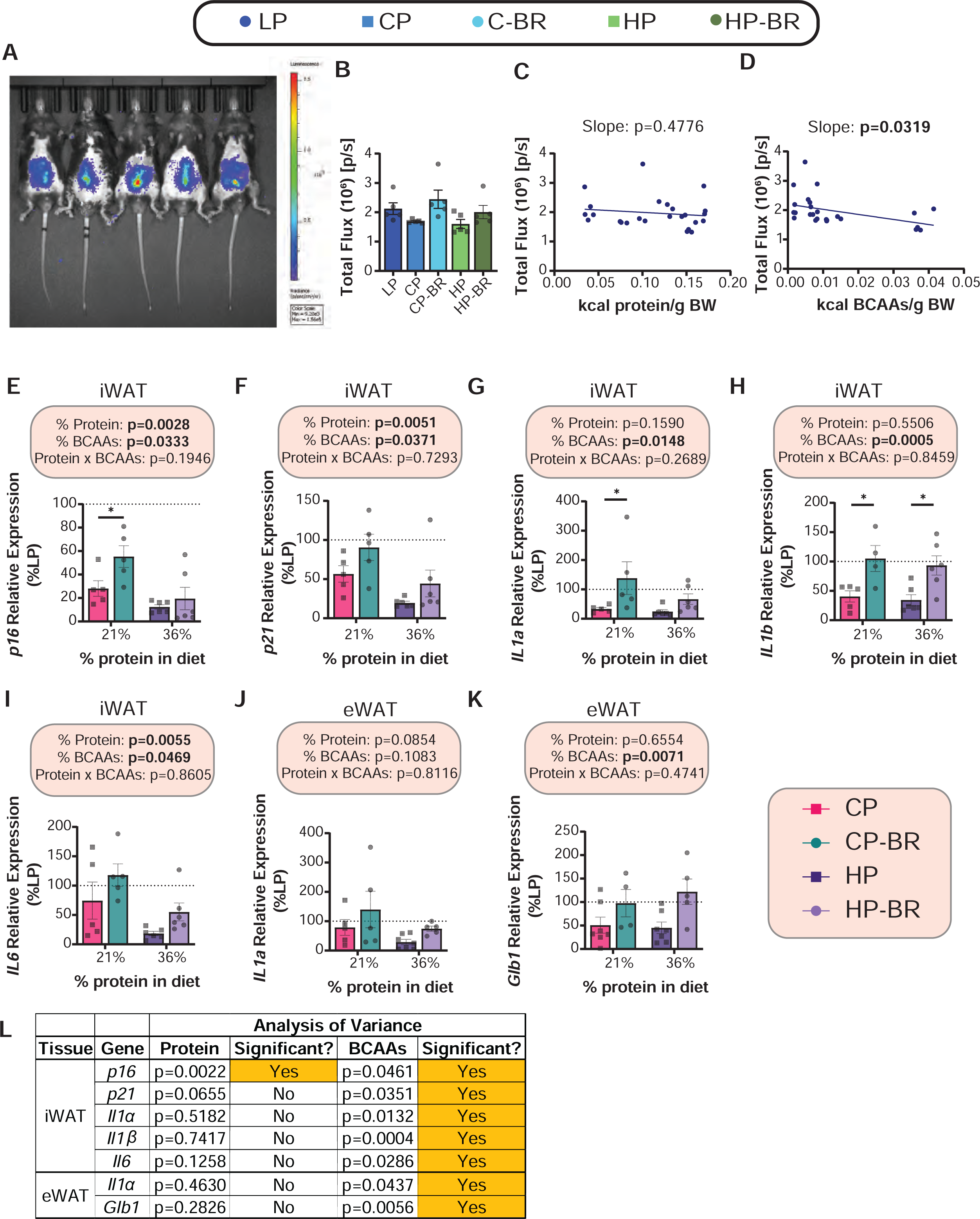
Diets low in BCAAs promote senescent cell accumulation in the adipose tissue of p16-3MR male mice. (A-B) *In vivo* bioluminescence of p16 after 22 weeks on the indicated diets (A); total flux was quantified (B). (C-D) Linear regression of p16 total flux versus kcals of protein (C) and kcals of BCAAs (D) consumed. (E-I) iWAT mRNA expression of the indicated genes. (J-K) eWAT mRNA expression of the indicated genes. (L) MLR analysis of significant adipose tissue senescence genes. (B) *p<0.05; Tukey test post ANOVA. (C-D) Linear regression of p16 total flux as a function of protein (O) or BCAAs (P) kilocalories; slope displayed to determine relationship. (E-K) n=4-6 mice/group (100% = average expression of the indicated gene in the iWAT or eWAT of LP-fed mice); the overall effect of protein, BCAAs, and the interaction represent the p-value from a two-way ANOVA; *p<0.05, Sidak’s test post 2-way ANOVA. (L) Statistics for the p-value are from a MLR analysis to determine the contribution of protein versus BCAAs from the senescence data set. Data represented as mean ± SEM.

We therefore decided to examine the adipose depots, specifically, the iWAT and epididymal white adipose tissue (eWAT) since they are likely to show bioluminescence in the abdominal area, where we detected strong p16 activity, and diets have been shown to have a robust effect on adipose tissue senescence in other studies ^62–65^. In contrast to our original hypothesis that BCAA restriction would reduce CS, in our original C57BL/6J mice we found a strong increase in the expression of multiple CS and SASP genes in BCAA-restricted mice, with the expression of *p16, p21, Il1a, Il1b,* and *Il6* increasing in BCAA-restricted iWAT, and *Il1a* and *Glb1* expressing in BCAA-restricted eWAT (**Figs. 2E-K**). This interpretation was further supported by multiple linear regression (MLR) analysis, which found a strong contribution of BCAA levels – but not protein level – to the expression of these CS and SASP genes (**Fig. 2L**). Broadly, we find that opposed to what we observed in the liver, BCAA restriction increases CS in adipose tissue, increasing expression of most CS and SASP genes in iWAT, increasing expression of *Il1a* and *Glb1* expression in eWAT, and increasing CS genes in BAT as well (**Table S3 and S4**).

The differential effect of BCAAs on CS in the liver and iWAT was quite surprising. We hypothesized that this might be due in part to the differential effect of FGF21, a hormone induced by protein restriction and BCAA restriction that has been shown to regulates CS ^46,47^. FGF21 has differential effects on mTOR protein kinase signaling in liver and adipose tissue ^43^, and mTOR signaling is a key regulator of aging, metabolism, and components of the CS and SASP ^18^. We examined *Fgf21* expression in the liver and three adipose depots; *Fgf21* expression was higher in the liver and iWAT of LP-fed mice than all other groups, with CP-BR-fed mice having the next highest expression; there were minimal effects of BCAAs on *Fgf21* expression in iWAT or BAT (**Supplementary Figs. S6A-D**). Therefore, we then looked at mTORC1 signaling in the liver and adipose tissue. There were minimal effects of diet on the phosphorylation of the mTORC1 substrates S6K1 T389 or 4E-BP1 T37/S46, or of the downstream readout S6 S240/S244 in either of the tissues (**Supplementary Figs. S6E-J**).

To gain insight into what types of damage might be signaling liver and iWAT cells to undergo CS, we looked at several genes related to different types of DNA damage. We found that a BCAA diet primarily affected the expression of genes related to mitochondrial dysfunction and mitochondrial biogenesis, without altering expression of genes related to antioxidant function and oncogenesis (**Figs. 3A-B**). Specifically, we found that the livers of CP-BR-fed mice had lower *Cycl1* and *Sdha* expression as well as increased *Pgc1a* expression compared to CP-fed mice, suggesting they had improved mitochondrial function (**Fig. 3A**). In contrast, we found the opposite gene expression changes in the iWAT of CP-BR-fed mice, suggesting they had impaired mitochondrial function (**Fig. 3B**).

**Figure 3:**
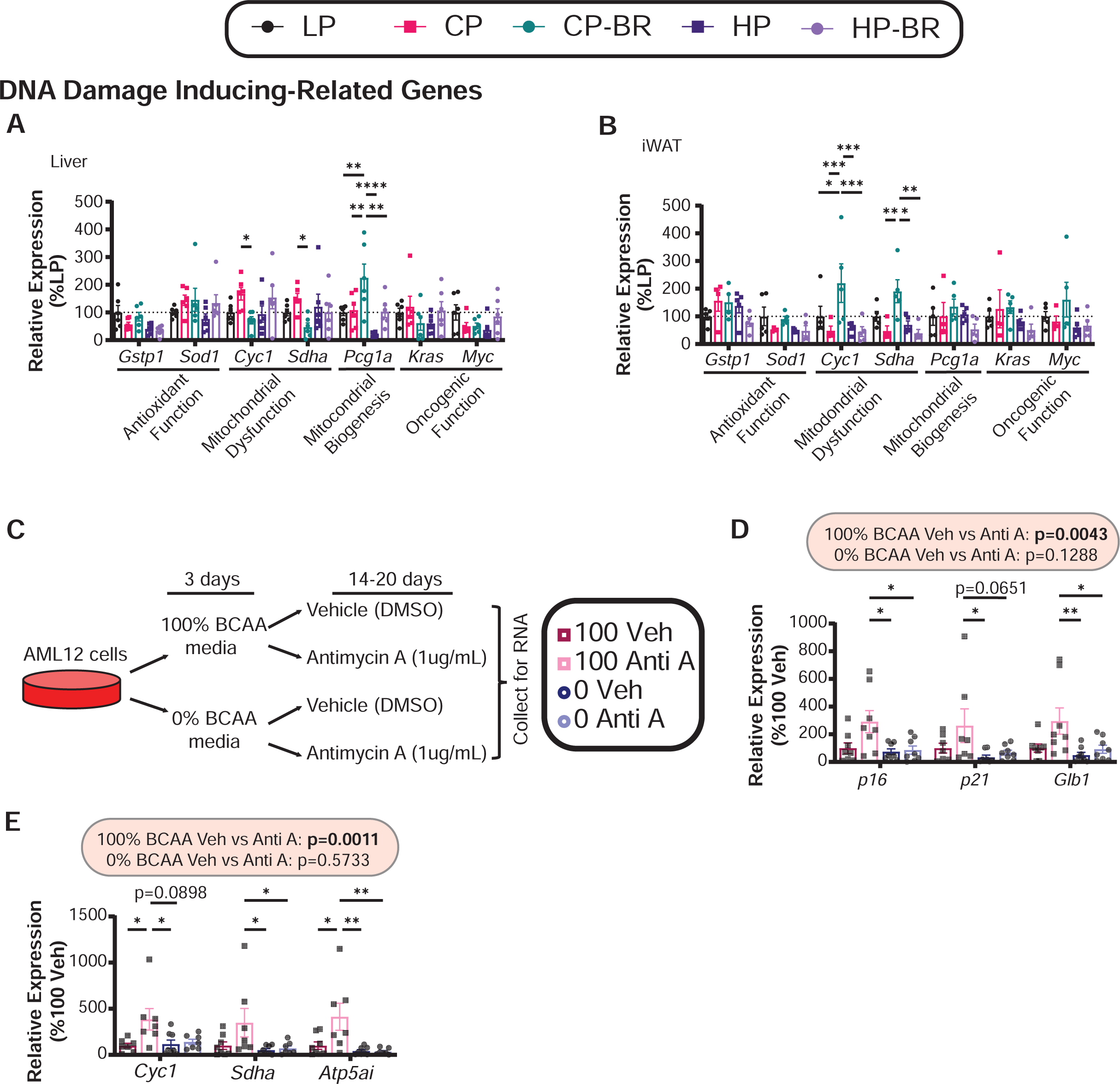
Lowering BCAAs protect from mitochondrial-dysfunction-related senescence in hepatocytes. (A-B) Hepatic (A) and iWAT (B) mRNA expression of DNA-damage-related genes. n=5-6 mice/group; *p<0.05, Tukey test post 2-way ANOVA conducted separately for each gene. (C) Experimental design for cell culture. (D-E) mRNA expression of *p16, p21* and *Glb1* (D) and mitochondrial-related genes (*Cyc1, Sdha* and *Atp5a1*) (E). n=8 biological replicates/group; the overall effect of Antimycin A (Anti A) from a 2-way ANOVA conducted separately for 100% BCAAs and 0% BCAAs; *p<0.05, Tukey test post 2-way ANOVA conducted separately for each gene. Data represented as mean ± SEM.

This suggested to us that BCAAs, specifically at 21% protein, may mediate mitochondrial function/dysfunction, leading to the tissue-specific differences in senescence. To confirm that mitochondrial dysfunction related to dietary BCAA may be mediating the effects on senescence, we turned to cell culture. We cultured mouse AML12 hepatocytes in media containing the normal level of BCAAs (100% BCAA) or restricted BCAAs (0% additive BCAA; this media still contains some BCAAs from serum). We treated these cells with antimycin A for 14-20 days to induce mitochondrial dysfunction-related CS, and then collected the cells to assess CS (**Fig. 3C**). As we expected, AML12 cells treated with antimycin A senesced, with increased expression of *p16, p21* and *Glb1* (**Fig. 3D**). Culture of cells in the BCAA-restricted media (0% added BCAA) prevented antimycin-A induced CS. This difference in CS was associated with antimycin A-induced mitochondrial dysfunction, which occurred in cells cultured in 100% BCAA and which cells cultured in 0% BCAA media were protected from (**Fig. 3E**). Additionally, it was recently shown that DNA-damage leads to altered mitochondrial structure and dynamics, which downregulates fatty acid oxidation and induces cellular senescence ^66^. Therefore, we looked at gene expression of lipolysis, the initial step of fatty acid oxidation, in both tissues and found that at least at 36% protein, lipolytic gene expression is increased in the liver and decreased in the iWAT (**Supplementary Figs. 6E-F**). These results support a model in which BCAA restriction protects liver cells from CS via cell-autonomous effects on mitochondrial dysfunction-related senescence induction.

### Restriction of each individual BCAA does not replicate the effects of restricting all three BCAAs on hepatic cellular senescence

The metabolic benefits of the CP-BR diet are due primarily to the restriction of isoleucine, with a lesser contribution from restriction of valine ^52^. To determine if the effects of CP-BR on cellular senescence is due to a single BCAA, we placed 12-week-old C57BL/6J male mice on the CP and CP-BR diets, as well on leucine-restricted (Leu-R), isoleucine-restricted (Ile-R) or valine-restricted (Val-R) diets. All of these diets are isocaloric with identical levels and sources of fats and carbohydrates, and are isonitrogenous through the addition of non-essential amino acids in restricted diets; the full diet composition is provided in **Table S1**.

Over the course of 17 weeks, we tracked physiological parameters in all groups of mice (**Fig. 4A**). As anticipated, CP-BR-, Ile-R- and Val-R-fed mice consumed more food than CP and Leu-R-fed mice, yet these groups showed attenuated weight and fat mass gain (**Figs. 4B-F and Supplementary Figs. 7A-D**) as a result of increased energy expenditure (**Supplementary Figs. 7F-H**). These phenotypes were associated with strong induction of FGF21 in the blood of CP-BR, Ile-R, and Val-R-fed mice and increased expression of the thermogenic genes *Bmp8b* and *Elovl3* in the BAT of CP-BR- and Val-R-fed mice (**Supplementary Fig 7I-J**), and with improved blood glucose control in the same animals as well as Ile-R-fed mice (**Supplementary Figs. 7L-R**).

**Figure 4:**
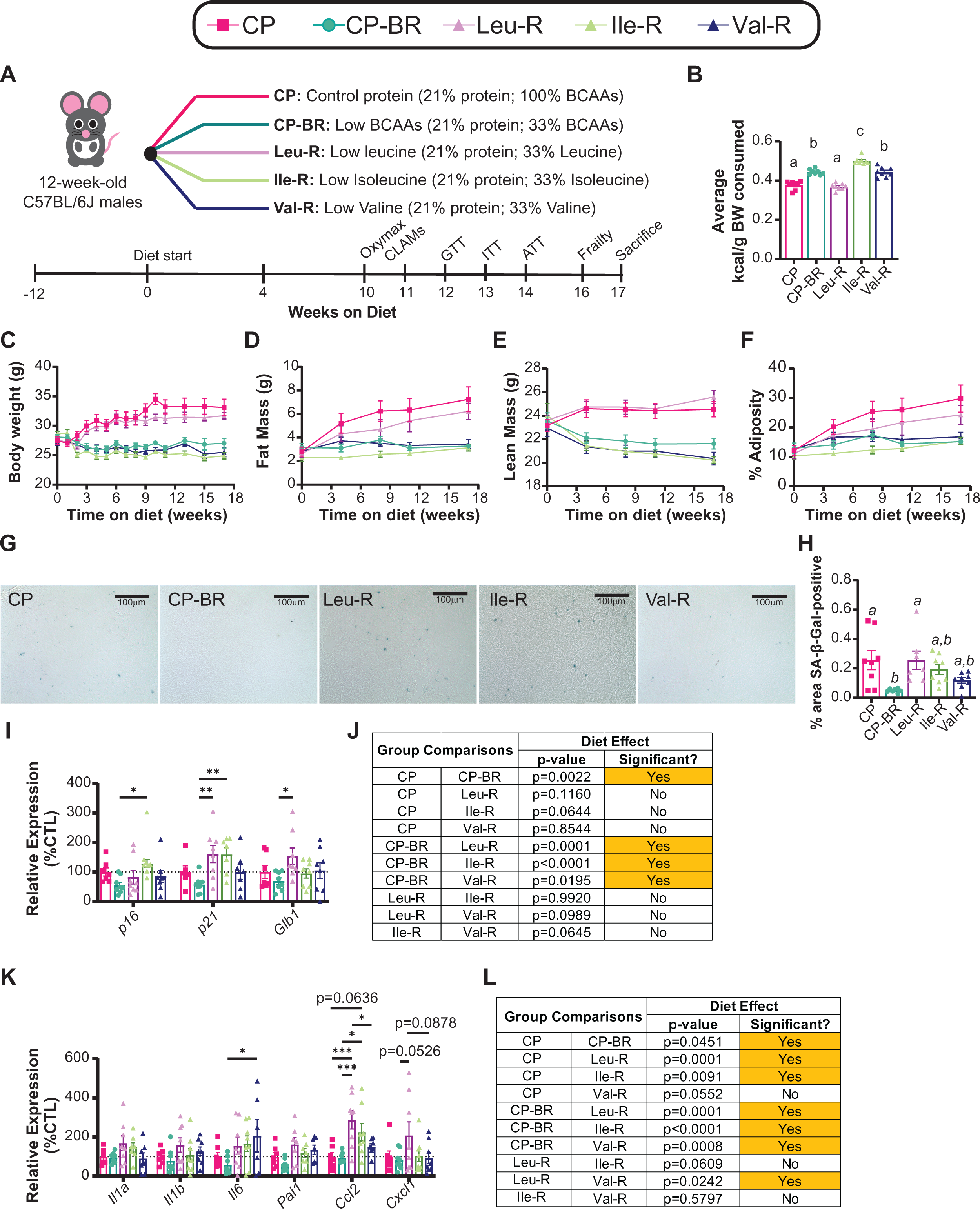
Restriction of each individual BCAA does not replicate the effects of a CP-BR diet on hepatic cellular senescence. (A) Experimental design. (B) Average consumption of kilocalories per gram of body weight (kcal/g BW) per group over 17-week period on diet. (C-F) Body weight (C), fat mass (D), lean mass (E), and adiposity (F) of mice fed the indicated diets over time. (G-H) Hepatic SA-βGal staining at 40X magnification (scale bar = 100 µm) (G) with quantification of SA-βGal-positive area (H). (I-J) Hepatic mRNA expression of *p16, p21* and *Glb1* (I) and the overall effect of each diet on senescence gene expression (J). (K-L) Hepatic SASP mRNA expression (K) and the overall effect of each diet on SASP gene expression (L). (B) n=7-8 mice/group; means with the same lowercase letter are not significantly different from each other, Tukey test post ANOVA, p<0.05.(H) n=7-8 mice/group; *p<0.05, Tukey test post ANOVA. (I-L) n=8 mice/group; the overall effect of diet from a 2-way ANOVA; *p<0.05, Tukey’s test post 2-way ANOVA conducted separately for each gene. Data represented as mean ± SEM.

Finally, we assessed hepatic senescence. Although CP-BR did reduce SA-βgal positivity and CS gene expression, Ile-R and Val-R did not despite their improvements in metabolic health (**Fig. 4G-L**). Interestingly, restriction of individual BCAAs seemed to increase, not decrease hepatic SASP expression (**Fig. 4K-L**). This suggests that while Ile-R and Val-R may mediate the metabolic and physiological effects seen in CP-BR diets, it does not mediate the effects of BCAA restriction on hepatic CS and the SASP.

### The effect of BCAAs on metabolism and CS is sex-specific

To examine the role of sex in the response of CS to dietary protein, we placed 14-week-old, female, p16-3MR mice on the five diets (LP, CP, CP-BR, HP, HP-BR) for 28 weeks (**Fig. 5A**). After 28 weeks on diet, we found largely no effect of diet in females (**Figs. 5B-C**). In agreement with this, BCAA-restricted females do not consume more food, but their consumption of protein and BCAAs was similar to the males (**Figs. 5D-F, Supplementary Figs. 8A-C, Supplementary Figs. 1M-P**). While females fed a low BCAA diet tended to have improved glucose tolerance (**Supplementary Fig. 8D-E**), BCAAs largely had no effect on fasting blood glucose, insulin sensitivity nor suppression of hepatic gluconeogenesis (**Supplementary Figs. 8F-J**).

**Figure 5:**
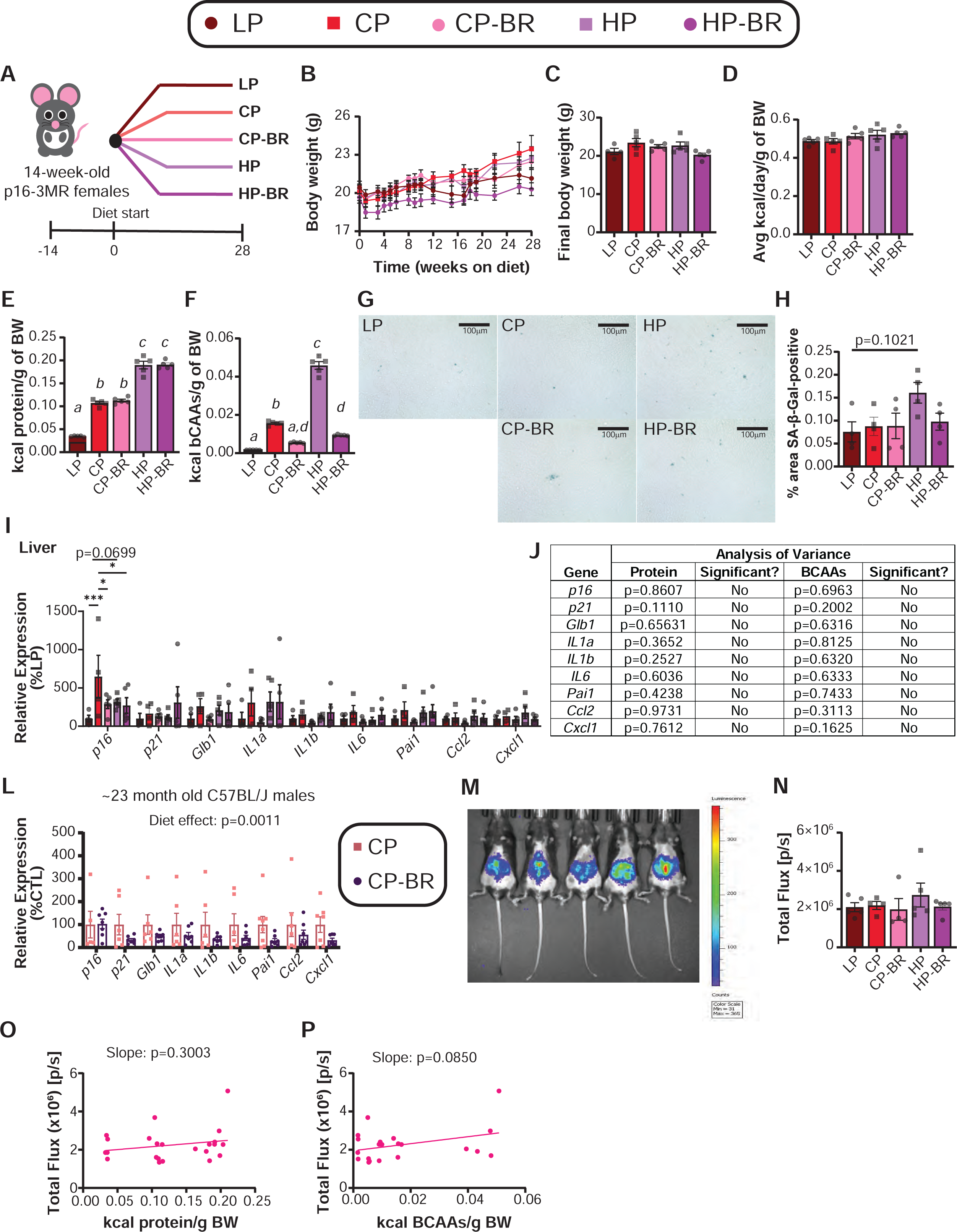
Low BCAA diets do not have the same benefits in female mice as they do in male mice. (A) Experimental design. (B-C) Body weight of mice fed the indicated diets over time (B) and final body weights (C). (D) Average kilocalories consumed per gram of body weight by each group of mice over the entire study. (E-F) Average kilocalories derived from protein (E) or derived from BCAAs (F) consumed by each group of mice over the entire study. (G-H) Hepatic SA-βGal staining at 40X magnification (scale bar = 100 µm) (G) with quantification of SA-βGal-positive area (H). (I-J) Hepatic mRNA expression of senescence and SASP genes (I) and MLR analysis to determine contribution of protein vs. BCAAs to gene expression (J). (L) Hepatic mRNA expression of senescence and SASP genes in female C57BL6/J mice on CP and CP-BR diets from 16 months of age until 23 months of age. (M-N) *In vivo* bioluminescence of p16 after 22 weeks on the indicated diets (M); total flux was quantified (N). (O-P) Linear regression of p16 total flux versus kcals of protein (O) and kcals of BCAAs (P) consumed. (B-J, M-P) n=4-5 female mice/group. (L) n=5-7 female mice/group. (C-F, H, N) means with the same lowercase letter are not significantly different from each other, Tukey test post ANOVA, p<0.05. (I) *p<0.05, Tukey test post 2-way ANOVA conducted separately for each gene. (J) Statistics for the p-value are from a MLR analysis to determine the contribution of protein versus BCAAs from the senescence data set. (I) *p<0.05, Sidak’s test post 2-way ANOVA conducted separately for each gene. (O-P) Linear regression of p16 total flux as a function of protein (O) or BCAAs (P) kilocalories; slope displayed to determine relationship. Data represented as mean ± SEM.

We examined CS in the livers of female p16-3MR mice. We found no significant difference in SA-βgal activity between any of the groups, though HP-fed females trended toward higher hepatic SA-βgal activity than LP-fed mice (p=0.1021) (**Figs. 5G-H**). While LP-fed and CP-BR-fed females did have reduced expression of *p16* compared to CP-fed females, this was the only significant difference identified; using the MLR approach we used previously, we found that neither dietary BCAAs nor dietary protein contributed to the expression of any of the CS and SASP genes we examined (**Figs. 5I-J**). While females are thought to be more prone to senescence ^67,68^, it may not be the case in young animals, such as in this study, as we see higher SA-βGal-positive staining in the liver of the males compared to females (**Figs. 1E-F**, **5G-H**). It may also be that in young mice, female-specific senescence accumulation is tissue-specific, and we would have seen CS in other tissues ^69^. As LP and CP-BR diets reduced *p16* expression in the liver (**Fig. 5I**), we decided to look at CS and SASP gene expression in the livers of C57BL/6J female mice fed a CP-BR diet for 6-7 months starting at 16 months of age until 23 months of age, using mRNA banked from a previous study ^29^. We found that there was an overall significant effect of diet on CS and SASP gene expression (**Fig. 5L**), suggesting that a reduction of dietary BCAAs can reduce hepatic senescence in middle-aged/aged C57BL/6J females.

We also assessed p16 bioluminsescence; we observed no significant differences in p16 expression as quantified by total flux in the females (**Figs. 5M-N**). While there was a trending positive relationship between flux and BCAAs consumed (p=0.085), we did not see significant effects of BCAAs when performing a MLR analysis in the iWAT (**Figs. 5O-P and Supplementary Figs. 8K-L**).

Overall, we found a sex-specific effect of BCAAs on metabolic health and senescence, with female mice largely not benefiting from BCAA restriction with respect to the effects on weight, body composition, and hepatic cellular senescence.

## Discussion

CS is one of the hallmarks of aging ^12^, and the long term accumulation of senescent cells has negative consequences on health and aging ^18^. Dietary interventions can protect against CS, with some studies showing that specific dietary macronutrients can impact CS ^18,30,63,65,70^. Dietary protein can promote hepatic senescence ^30^, and recent studies have shown that impaired catabolism of the BCAAs promotes CS ^36,37^.

Here, we have examined how dietary BCAAs and protein impact both metabolic health and CS. We find that diets low in BCAAs (LP, CP-BR and HP-BR) normalize body weight, preventing body weight gain and adiposity accretion, regardless of the overall level of dietary protein, in male mice. This effect is mediated by changes in energy balance, with restriction of BCAAs boosting food intake as well as energy expenditure via increased thermogenesis in the iWAT of LP-fed mice and in the BAT of CP-BR-fed and HP-BR-fed male mice. Male mice consuming diets with reduced levels of BCAAs also have improved glucose tolerance. Female mice generally have a substantially blunted response to reduction of BCAAs.

We see strong effects of BCAA restriction on CS in the liver, with a reduction in SA-βgal staining as well as CS and SASP gene expression, with robust effect in males under a variety of diets and ages as well as in aged females. Notably, our findings show that dietary protein content itself is not otherwise associated with hepatic CS or SASP gene expression, except insofar as higher protein diets normally contain higher levels of BCAAs. However, restricting BCAAs in the context of a HP diet does not protect against CS to the same extent as it does in the CP diet, suggesting that other components of the overall dietary context – for example, the decreased levels of carbohydrates in the HP diet – may impact the effect of BCAAs on hepatic CS. As the metabolic benefits of lower BCAAs are not lower in HP-fed mice, the benefits of BCAA restriction for CS are uncoupled from its effects on metabolic health.

The benefits of lower levels of BCAAs on CS in the liver are cell-autonomous, with BCAA restriction protecting the AML12 mouse hepatocyte cell line from mitochondrial dysfunction-induced senescence. However, the benefits are also cell-specific, with lower dietary BCAAs promoting CS in iWAT. Additional research will be required to fully understand both how lower levels of BCAAs protect against hepatic CS, how dietary context influences the effects of BCAAs, and why lower BCAAs potentiate CS in the iWAT without impairing metabolic health. It may be that there is some protective impact of CS in fat. Specifically, CS is important for wound healing ^71^, and perhaps the fat depot is in a state of healing from metabolic health. It will also be informative to identify effects of dietary BCAAs on CS in other tissues and cell types, and understand how sex interacts with BCAAs to influence CS. Finally, we were surprised to find that restriction of individual BCAAs did not influence CS, suggesting that simultaneously restriction of multiple BCAAs is required to reduce hepatic CS.

Limitations of our study include that these studies were primarily conducted in young, male mice. We chose to conduct the study in this way based on the results of a previous study of protein consumption and CS ^30^, but more pronounced differences may have been observed if we had used older mice with a higher spontaneous rate of CS or placed the mice on these diets for a longer period of time. Our mice were primarily sacrificed in the refed state following an overnight fast, and assessing tissues in a different feeding state could also provide additional insights. These studies were conducted exclusively in C57BL6/J mice, and we have shown that strain as well as sex contribute to the metabolic response to PR ^72^. Future studies should test the robustness of our results in other mouse strains as well as in genetically heterogenous mice.

In summary, recent work from multiple labs has shown that calorie quality, not just the total quantity, is a critical determinant of biological health, and that dietary protein in particular is a critical mediator of healthy aging ^19,20^. We and others have shown that many of the beneficial effects of low protein diets are mediated by reduced levels of the BCAAs, and that dietary levels of BCAAs are negatively associated with lifespan ^29,32^. Here, we find that reducing dietary BCAAs reduces hepatic CS via a cell-autonomous effect that is uncoupled from the beneficial effects of BCAA restriction on metabolic health. Surprisingly the effects of BCAA restriction are tissue-specific, and while low dietary BCAAs reduce hepatic CS, it increases CS in adipose tissue. Overall, these data support the notion that dietary composition is a critical regulator of CS burden during aging, and highlight BCAAs as the critical mediator of the effects of dietary protein on CS.

## Acknowledgements

We would like to thank all the members of the Lamming lab for their contributions to this work. We would like to thank Dr. Nicole Richardson for supplying the mRNA from middle-aged mice fed a BCAA restricted diet, and Justin Jefferies for assistance in performing the bioluminescence assays on the p16-3MR mice. We would also like to thank Dr. Matthew Yousefzadeh for his help troubleshooting the SA-β-Gal staining. The Lamming lab is supported in part by the NIH/NIA (AG056771, AG081482, AG084156, AG085898), the NIDDK (DK125859), and startup funds from UW-Madison. MFC is supported by F31AG082504. RB is supported by F31AG081115. MMS was supported in part by a Supplement to Promote Diversity in Health-Related Research RF1AG056771-06S1. DAH is a Wisconsin Alzheimer’s Disease Research Center REC scholar. The authors would like to acknowledge the Cancer Center Support Grant: NCI P30 CA014520, University of Wisconsin Small Animal Imaging & Radiotherapy Facility. The Lamming lab was supported in part by the U.S. Department of Veterans Affairs (I01-BX004031 and IS1-BX005524), and this work was supported using facilities and resources from the William S. Middleton Memorial Veterans Hospital. The content is solely the responsibility of the authors and does not necessarily represent the official views of the NIH. This work does not represent the views of the Department of Veterans Affairs or the United States Government

## Competing Interests

D.W.L. has received funding from, and is a scientific advisory board member of, Aeovian Pharmaceuticals, which seeks to develop novel, selective mTOR inhibitors for the treatment of various diseases.

## Author contributions

Experiments were performed in the Lamming laboratory. MFC, ARK, DAH, and DWL conceived the experiments and secured funding. MFC, IA, IDG, SML, PL, LEB, RB, MMS, JAI, BAK, and FX performed the experiments. MFC, IA, PL, LEB, SML, DM, and DWL analyzed the data. MFC, ARK, DAH and DWL wrote and edited the manuscript.

## Data Availability

The data that support the plots within this article and other findings of this study are provided as Source Data files.

