## Supplementary Figures and Supplementary Table Legends for "Tissue-specific effects of dietary protein on cellular senescence are mediated by branched-chain amino acids"

### Supplementary Figure 1

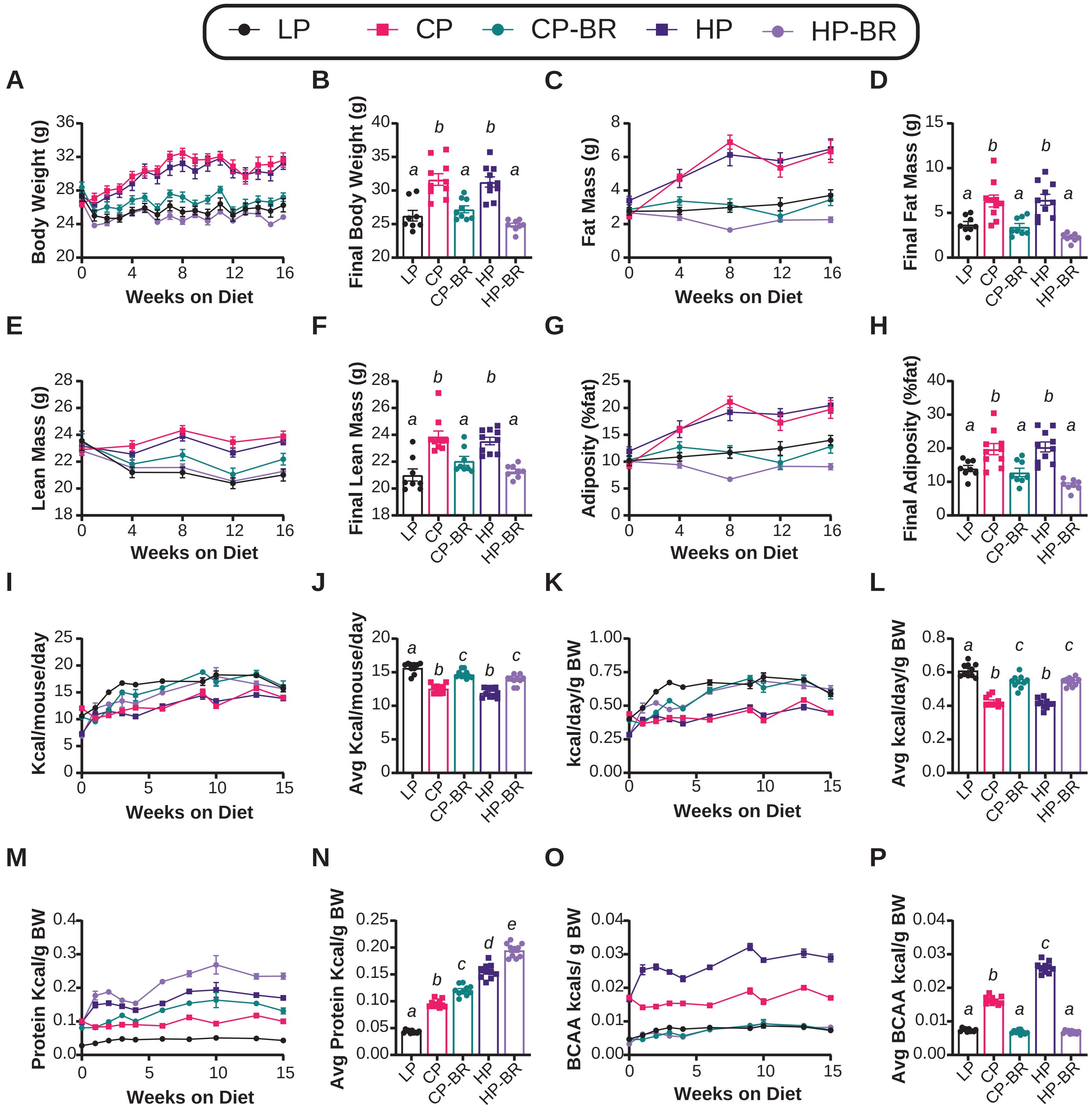

**Supplementary Figures**

**Supplementary Figure 1: Diets low in BCAAs normalize body weight without malnutrition.**

(A-B) Body weight of mice fed the indicated diets over time (A) and final body weight at end of study (B). (C-D) Fat mass over time (C) and final fat mass (D). (E-F) Lean mass over time (E) and final lean mass (F). (G-H) Percent adiposity over time (G) and final percent adiposity (H). (I-J) Kilocalories consumed over time (I) and average kilocalories consumed over the course of the entire study (J). (K-L) Kilocalories consumed per gram of body weight over time (K) and average kilocalories per gram of body weight consumed over the course of the entire study (L). (M-N) Kilocalories per gram of body weight derived from protein over time (M) and average kilocalories per gram of body weight consumed over the course of the entire study (N). (O-P) Kilocalories per gram of body weight derived from BCAAs over time (O) and average kilocalories per gram of body weight consumed over the course of the entire study (P). (A-P) n=10 mice/group. (B, D, F, H, J, L, N, P) means with the same lowercase letter are not significantly different from each other, Tukey test post ANOVA, $p<0.05$. Data represented as mean ± SEM.

### Supplementary Figure 2

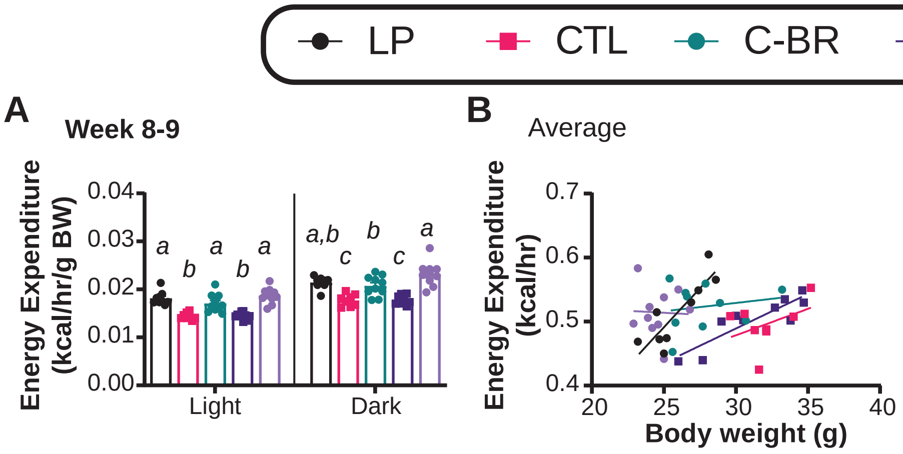


C

| Diet | | Slope | | Elevation | |
|---|---|---|---|---|---|
| | | p-value | Significant? | p-value | Significant? |
| LP | CTL | 0.1016 | No | 0.0028 | Yes |
| LP | C-BR | 0.0145 | Yes | Cannot be determined | |
| LP | HP | 0.0207 | Yes | Cannot be determined | |
| LP | HP-BR | 0.0504 | No | 0.28 | No |
| CTL | C-BR | 0.5456 | No | 0.0786 | No |
| CTL | HP | 0.7003 | No | 0.2446 | No |
| CTL | HP-BR | 0.0492 | No | 0.3010 | No |
| C-BR | HP | 0.1276 | No | 0.0118 | Yes |
| C-BR | HP-BR | 0.7552 | No | 0.8954 | No |
| HP | HP-BR | 0.2206 | No | 0.0126 | Yes |

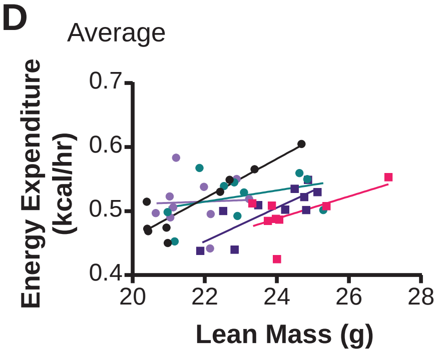


E

| Diet | | Slope | | Elevation | |
|---|---|---|---|---|---|
| | | p-value | Significant? | p-value | Significant? |
| LP | CTL | 0.2221 | No | 0.0004 | Yes |
| LP | C-BR | 0.0383 | Yes | Cannot be determined | |
| LP | HP | 0.6132 | No | p<0.0001 | Yes |
| LP | HP-BR | 0.0747 | No | 0.9261 | No |
| CTL | C-BR | 0.0517 | No | 0.0251 | Yes |
| CTL | HP | 0.4452 | No | 0.1706 | No |
| CTL | HP-BR | 0.4139 | No | 0.1145 | No |
| C-BR | HP | 0.1296 | No | 0.0326 | Yes |
| C-BR | HP-BR | 0.6933 | No | 0.9942 | No |
| HP | HP-BR | 0.1462 | No | 0.0467 | Yes |

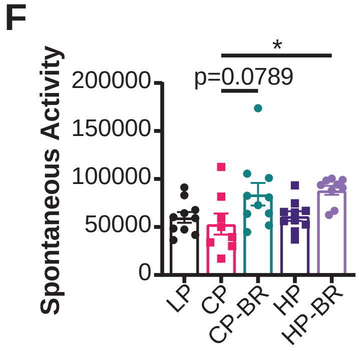


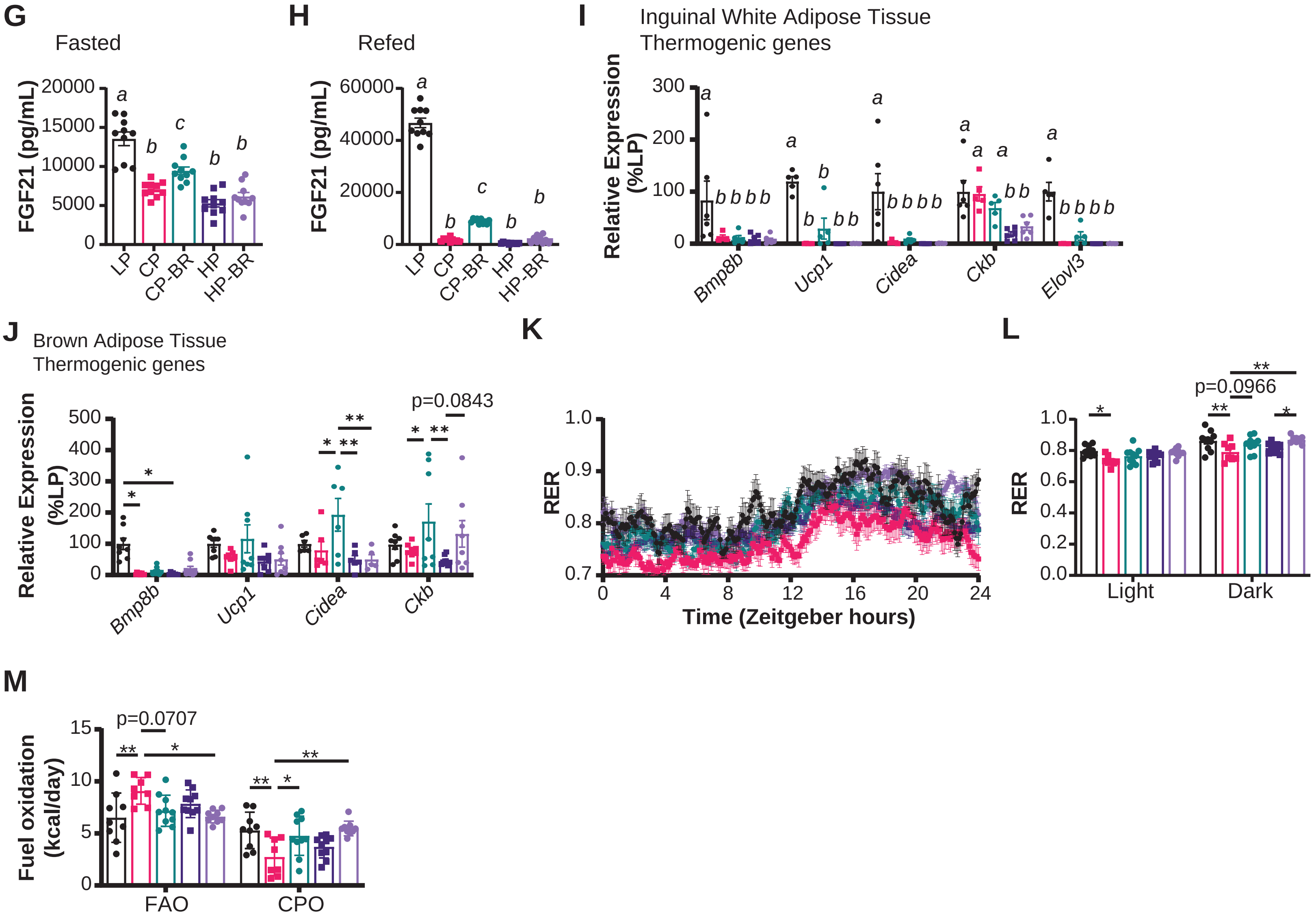

**Supplementary Figure 2: Restriction of BCAAs increases energy expenditure via brown adipose tissue thermogenesis.**

(A) Average light and dark energy expenditure. (B-E) Average energy expenditure over 24-hour period as a function of body weight (B-C) or lean mass (D-E). (F) Total spontaneous activity over 24-hour period per group. (G-H) Circulating FGF21 in the fasted (G) and refed (H) state. (I-J) mRNA expression of thermogenic genes in iWAT (I) and BAT (J). (K-L) Respiratory exchange ratio (RER) over 24-hour period represented in Zeitgeber hours (K) and their averaged values (L). (M) Fuel oxidation of fats and carbohydrates or protein. (A-H, K-M) n=8-10 mice/group; collected during weeks 8-9.(A) means with the same lowercase letter are not significantly different from each other, Tukey test post ANOVA conducted separately for the light and dark cycles, $p<0.05$. (C, E) simple linear regression (analysis of covariance) was calculated to determine if the slopes or elevations are equal; if the slopes are significantly different, differences in elevation cannot be determined. (F) *$p<0.05$, Tukey test post ANOVA. (G-H) means with the same lowercase letter are not significantly different from each other, Tukey test post ANOVA, $p<0.05$. (I) n=5-6 mice/group; means with the same lowercase letter are not significantly different from each other, Tukey test post 2-way ANOVA conducted separately for each gene, $p<0.05$. (J) n=7-8 mice/group; *$p<0.05$, Tukey test post 2-way ANOVA conducted separately for each gene. (L-M) *$p<0.05$, Tukey test post ANOVA conducted separately for the light and dark cycles (L) and for FAO/CPO (M). Data represented as mean ± SEM.

#### Supplementary Figure 3

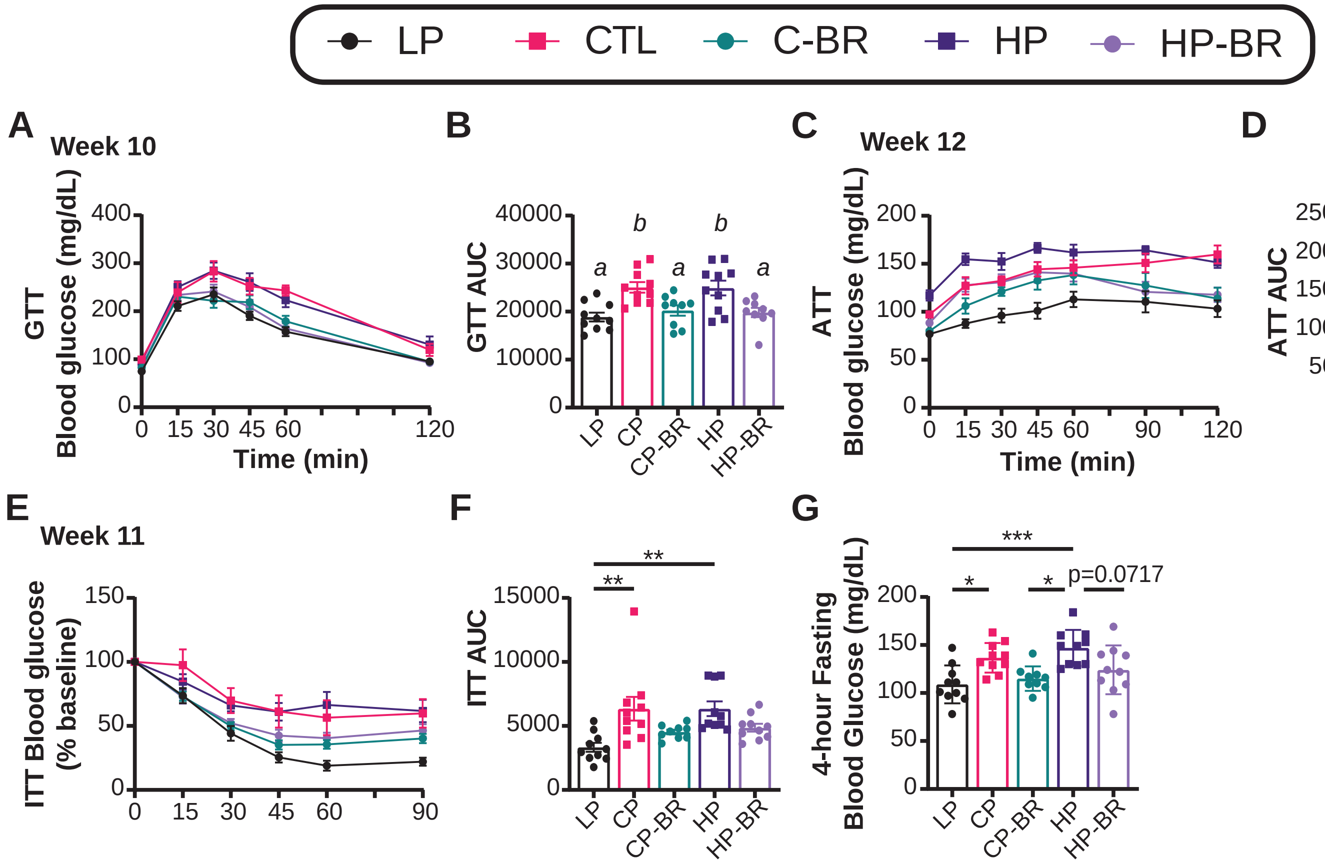

**Supplementary Figure 3: Diets low in BCAAs improve glucose homeostasis.**

(A-B) Glucose tolerance test conducted after 10 weeks on diet (A) and quantified area under the curve (B). (C-D) Suppression of hepatic gluconeogenesis as assessed by alanine tolerance test conducted after 12 weeks on diet (C) and quantified area under the curve (D). (E-F) Insulin tolerance test conducted after 11 weeks on diet (E) and quantified area under the curve (F). (G) 4-hour fasting blood glucose levels were collected after 11 weeks on diet. (A-G) n=10 mice/group. (B, D) means with the same lowercase letter are not significantly different from each other, Tukey test post ANOVA, $p<0.05$. (F, G)*$p<0.05$, Tukey test post ANOVA. Data represented as mean ± SEM.

### Supplementary Figure 4

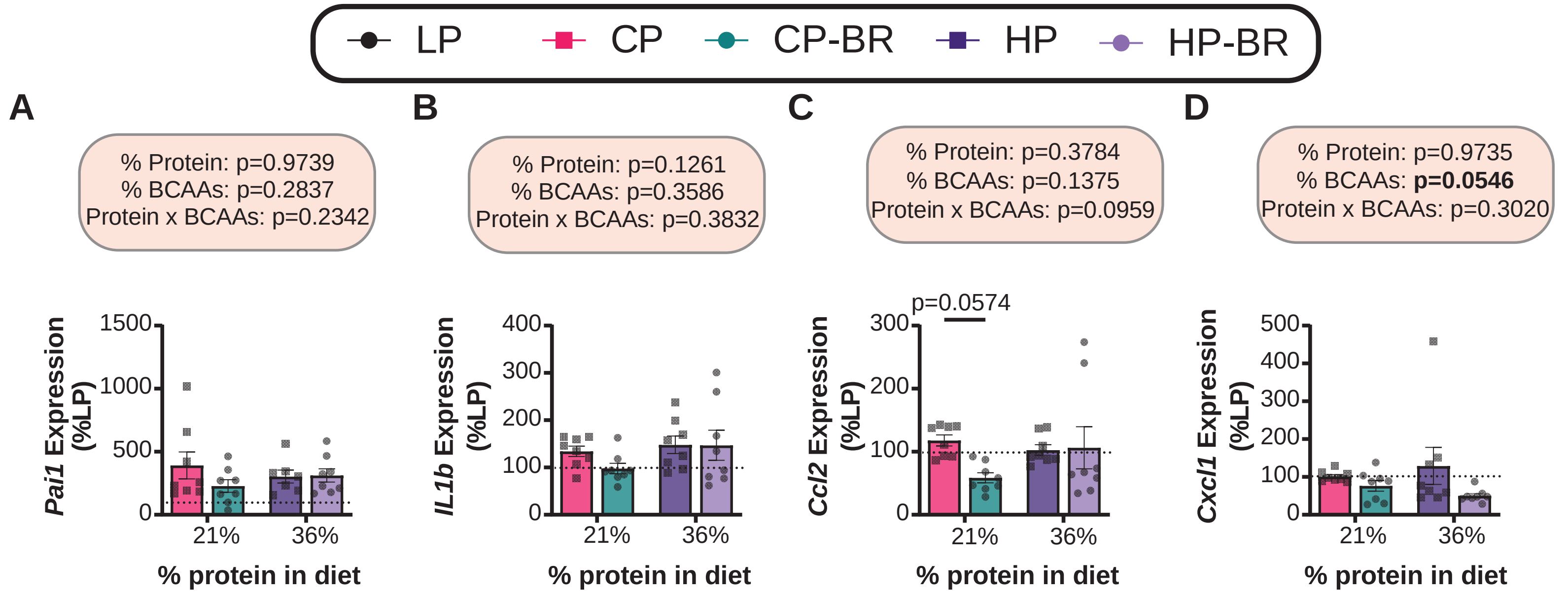

**Supplementary Figure 4: Hepatic SASP mRNA expression of male mice.**

(A-D) Hepatic mRNA expression of *Pai1, IL1b, Ccl2* and *Cxcl1.* n=8 mice/group. 100% = average expression of the indicated gene in the liver of LP-fed mice. The overall effect of protein, BCAAs, and the interaction represent the p-value from a two-way ANOVA; $*p<0.05$, Sidak's test post 2-way ANOVA. Data represented as mean ± SEM.

### Supplementary Figure 5

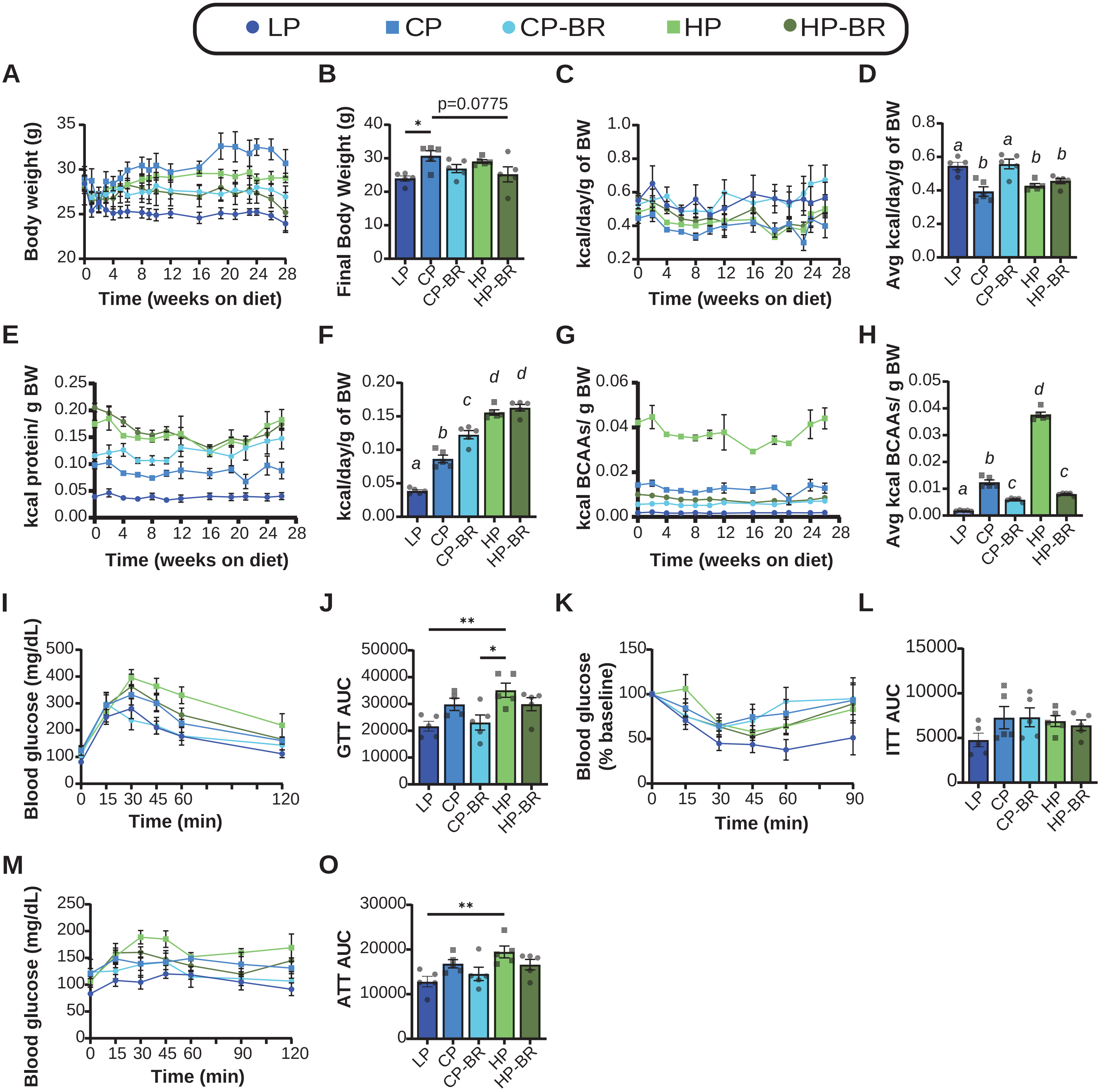

**Supplementary Figure 5: Diets low in BCAAs have similar benefits on metabolic health in p16-3MR male mice.**

(A-B) Body weight over 28-week period per group (A) and final body weight (B). (C-D) Kilocalories per gram of body weight consumed over time (C) and averaged over entire experiment (D). (E-F) Average kilocalories derived from protein (E) and kilocalories derived from BCAAs (F) consumed over entire study. (G-H) Glucose tolerance test conducted after 12 weeks on diet (G) and quantified area under the curve (H). (I-J) Insulin tolerance test conducted after 13 weeks on diet (I) and quantified area under the curve (J). (K-L) Alanine tolerance test conducted after 14 weeks on diet (K) and quantified area under the curve (L). (A-O) n=4-5 male mice/group. (B, J, L, O) *$p<0.05$, Tukey test post ANOVA. (D, F, H) means with the same lowercase letter are not significantly different from each other, Tukey test post ANOVA, $p<0.05$. (B, D, F, H, J, L, O) Data for each individual mouse is plotted; statistics for the p-value from a one-way ANOVA. Data represented as mean ± SEM.

### Supplementary Figure 6

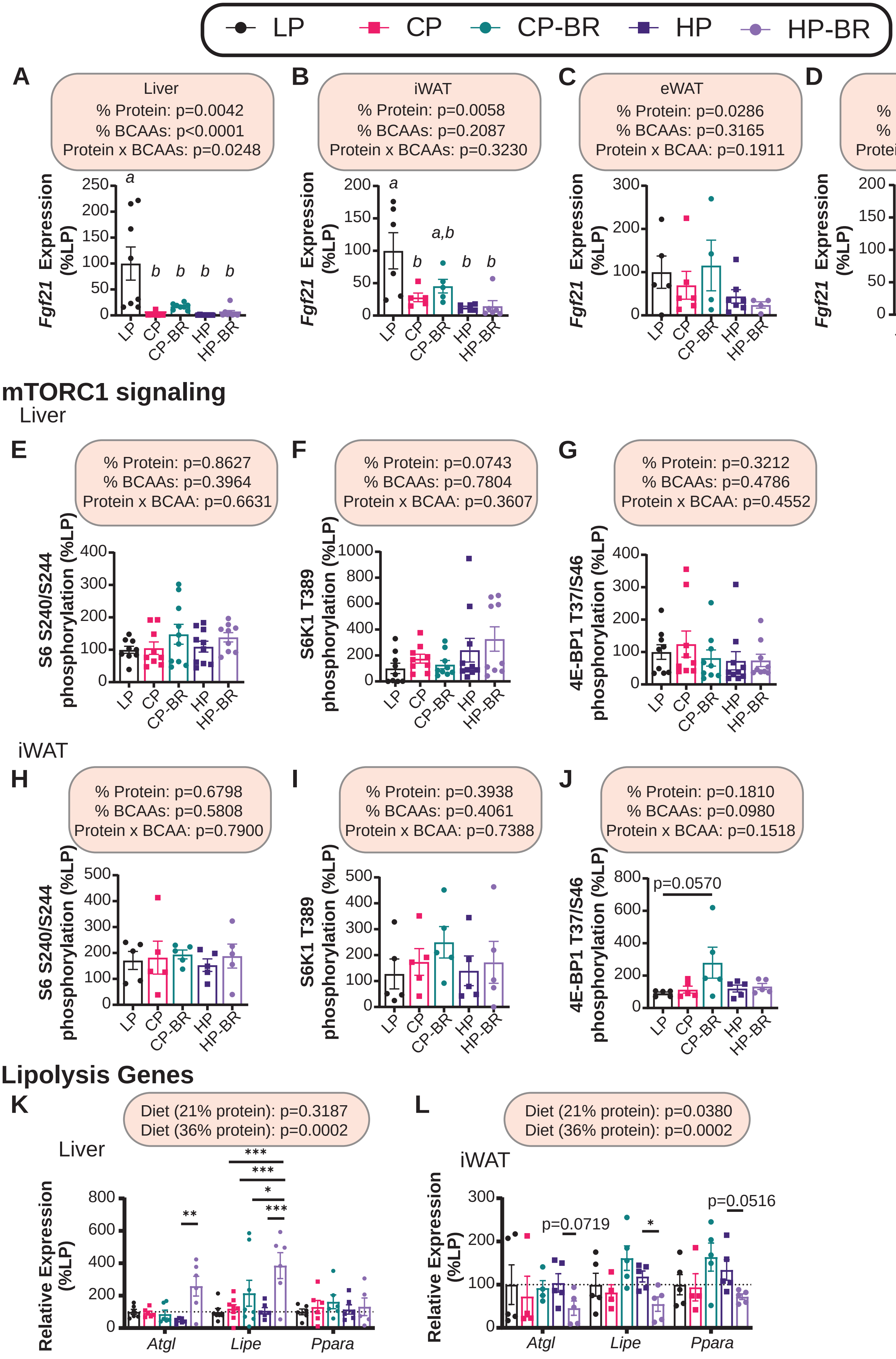

**Supplementary Figure 6: Low BCAA diets do not induce a tissue-specific effect on the FGF21/mTORC1 signaling pathway.**

(A-D) mRNA expression of *Fgf21* in liver (A), iWAT (B), eWAT (C) and BAT (D). (E-J) Phosphorylation of S6 S240/S244 (E, H), S6K1 T389 (F, I) and 4E-BP1 T37/S46 (G, J) in liver (E-G) and iWAT (H-J) via western blot. (K-L) mRNA expression of lipolysis genes in liver (K) and iWAT (L). (A-J) n=4-8 mice/group. The overall effect of protein, BCAAs, and the interaction represent the p-value from a two-way ANOVA for the CP, CP-BR, HP and HP-BR groups only. means with the same lowercase letter are not significantly different from each other, Tukey test post 1-way ANOVA, $p<0.05$. (K-L) n=4-6 mice/group; the overall effect of dietary BCAAs in the context of control protein (21%) and high protein (36%) from a 2-way ANOVA conducted separately for 100% BCAAs and 0% BCAAs; *$p<0.05$, Tukey test post 2-way ANOVA conducted separately for each gene Data represented as mean ± SEM.

### Supplementary Figure 7

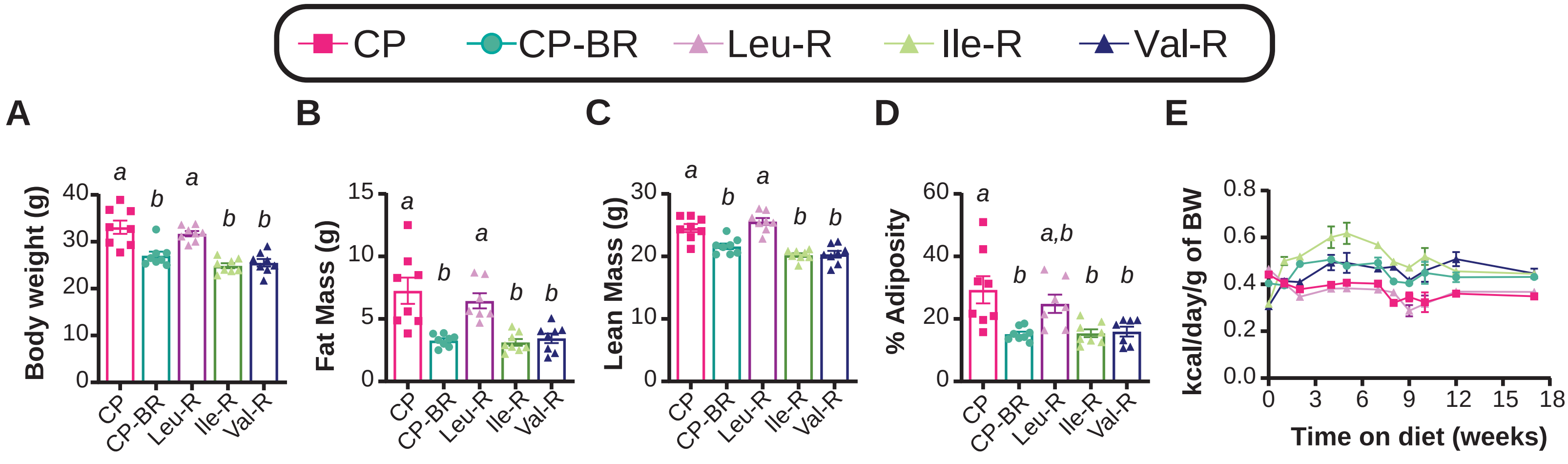


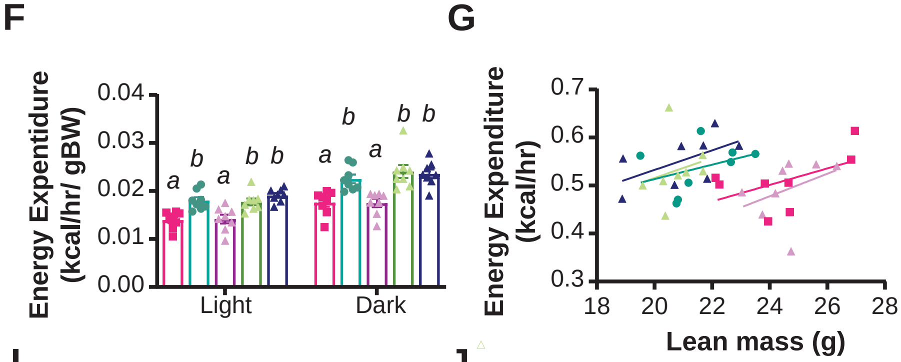


H

| Diet | | Slope | | Elevation | |
|---|---|---|---|---|---|
| | | p-value | Significant? | p-value | Significant? |
| CTL | C-BR | 0.9106 | No | 0.0484 | Yes |
| CTL | Leu-R | 0.8084 | No | 0.4954 | No |
| CTL | Ile-R | 0.9186 | No | 0.1077 | No |
| CTL | Val-R | 0.8515 | No | 0.0106 | Yes |
| C-BR | Leu-R | 0.7450 | No | 0.0054 | Yes |
| C-BR | Ile-R | 0.8761 | No | 0.8079 | No |
| C-BR | Val-R | 0.7853 | No | 0.3040 | No |
| Leu-R | Ile-R | 0.9518 | No | 0.1368 | No |
| Leu-R | Val-R | 0.9064 | No | 0.0145 | Yes |
| Ile-R | Val-R | 0.9904 | No | 0.5661 | No |

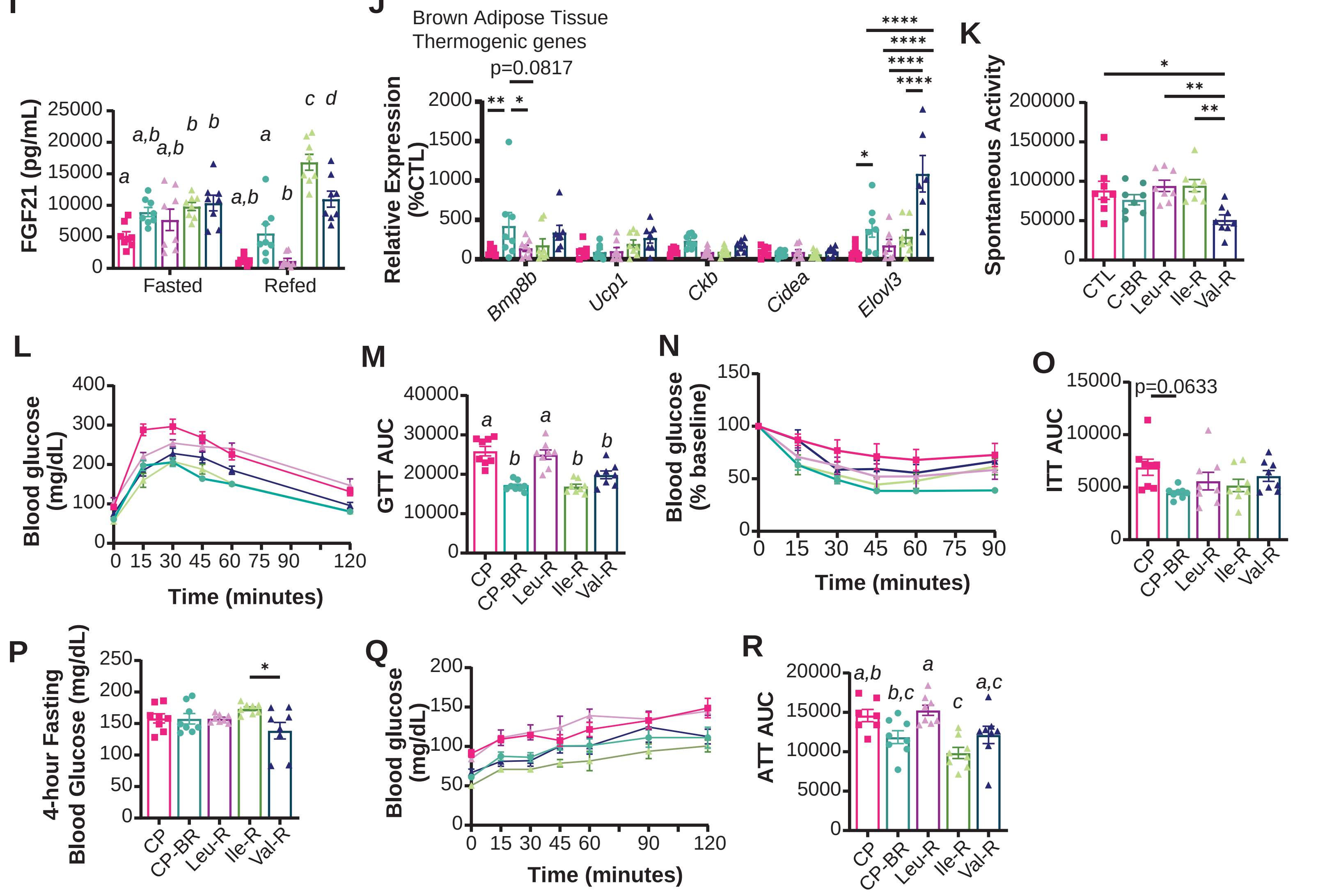

**Supplementary Figure 7: The benefits of CP-BR on the metabolic health of C57BL/6J males are mediated by the restriction of isoleucine and valine.**

(A-D) Final body weight (A), fat mass (B), lean mass (C), and percent adiposity (D) after 17 weeks on diet. (E) Kilocalories per day per gram of body weight. (F) Energy expenditure per gram of body weight in the light and dark phase. (G-H) Averaged energy expenditure over 24-hour period as a function of lean mass. (I) Circulating FGF21 in the fasted and refed state. (J) mRNA expression of thermogenic genes in BAT. (K) Total spontaneous activity over 24-hour period. (L-M) Glucose tolerance test conducted after 12 weeks on diet (L) and quantified area under the curve (M). (N-O) Insulin tolerance test conducted after 13 weeks on diet (N) and quantified area under the curve (O). (P) 4-hour fasting blood glucose levels taken after 13 weeks on diet. (Q-R) Suppression of hepatic gluconeogenesis as assessed by alanine tolerance test conducted after 14 weeks on diet (Q) and quantified area under the curve (R). (A-R) n=8 mice/group. (A-D, M, R) means with the same lowercase letter are not significantly different from each other, Tukey test post ANOVA, $p<0.05$. (F, I) means with the same lowercase letter are not significantly different from each other, Tukey test post ANOVA conducted separately for the light and dark cycles, $p<0.05$. (H) simple linear regression (analysis of covariance) was calculated to determine if the slopes or elevations are equal; if the slopes are significantly different, differences in elevation cannot be determined. (J) *$p<0.05$, Tukey test post 2-way ANOVA conducted separately for each gene. (K, O-P) *$p<0.05$, Tukey test post ANOVA, Data represented as mean ± SEM.

### Supplementary Figure 8

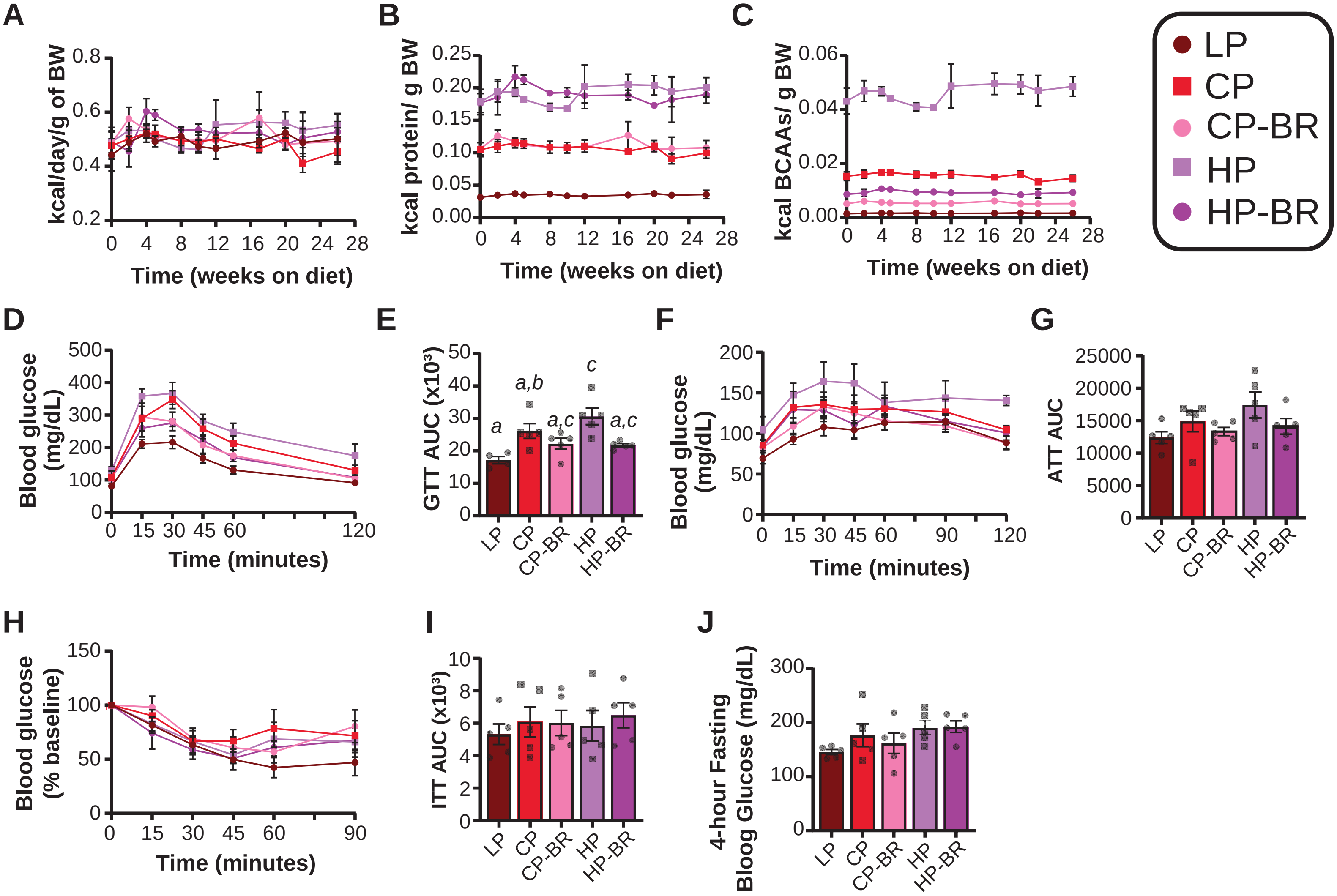


K

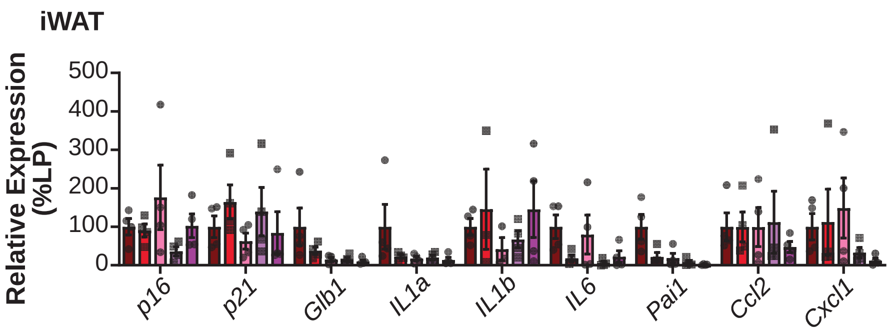


L

| Gene | Analysis of Variance | | | |
|---|---|---|---|---|
| | Protein | Significant? | BCAAs | Significant? |
| *p16* | p=0.6990 | No | p=0.1225 | No |
| *p21* | p=0.4536 | No | p=0.2153 | No |
| *Glb1* | p=0.0180 | Yes | p=0.5754 | No |
| *IL1a* | p=0.0671 | No | p=0.6970 | No |
| *IL1b* | p=0.7492 | No | p=0.9931 | No |
| *IL6* | p=0.2837 | No | p=0.8403 | No |
| *Pai1* | p=0.0037 | Yes | p=0.6156 | No |
| *Ccl2* | p=0.4387 | No | p=0.3945 | No |
| *Cxcl1* | p=0.3943 | No | p=0.9792 | No |

**Supplementary Figure 8: Diets low in BCAAs largely do not impact metabolic health in female mice**

(A) Kilocalories per gram of body weight consumed over time. (B-C) Kilocalories consumed derived from protein (B) and BCAAs (C) over time. (D-E) Glucose tolerance test conducted after 12 weeks on diet (D) and quantified area under the curve (E). (F-G) Suppression of hepatic gluconeogenesis as assessed by alanine tolerance test conducted after 14 weeks on diet (F) and quantified area under the curve (G). (H-I) Insulin tolerance test conducted after 13 weeks on diet (H) and quantified area under the curve (I). (J) 4-hour fasting blood glucose levels taken after 13 weeks on diet. (K-L) mRNA expression of senescence genes in iWAT (K) with the statistics of contribution of protein and BCAAs to their expression (L). (A-L) n=4-5 female mice/group. (E, G, I-J) means with the same lowercase letter are not significantly different from each other, Tukey test post ANOVA, $p<0.05$. (K) *$p<0.05$, Tukey test post 2-way ANOVA conducted separately for each gene. (L) Statistics for the p-value are from a MLR analysis to determine the contribution of protein versus BCAAs from the senescence data set. Data represented as mean ± SEM.

**Supplementary Table Legends**

**Supplementary Table 1: Experimental Diets**

The composition and calorie content of the experimental diets used in this study.

**Supplementary Table 2: qRT-PCR primer sequences**

Forward and reverse primer sequences for qRT-PCR.

**Supplementary Table 3: Interaction, gene effect and diet effect in the liver, iWAT, eWAT, and BAT on senescence gene expression.**

Values between indicated groups represent the p-value from a two-way ANOVA; *$p<0.05$, Sidak's test post 2-way ANOVA. n=5-8 mice/group.

**Supplementary Table 4: Multiple linear regression analysis values from iWAT, eWAT and BAT.**

MLR analysis p-value for the contribution of protein versus BCAAs in three adipose depots. n=5-8 mice/group.

**Supplementary Table 5: Cell culture media composition.**

Cell culture media components and their catalog numbers.
